# Cerebellar nuclei neurons dictate cortical growth through developmental scaling of presynaptic Purkinje cells

**DOI:** 10.1101/310953

**Authors:** Ryan T. Willett, Alexandre Wojcinski, N. Sumru Bayin, Zhimin Lao, Daniel Stephen, Katherine L. Dauber-Decker, Zhuhao Wu, Marc Tessier-Lavigne, Hiromitsu Saito, Noboru Suzuki, Alexandra L. Joyner

## Abstract

Efficient function of neural systems requires the production of specific cell types in the correct proportions. Here we report that reduction of the earliest born neurons of the cerebellum, excitatory cerebellar nuclei neurons (eCN), results in a subsequent reduction in growth of the cerebellar cortex due to an accompanying loss of their presynaptic target Purkinje cells. Conditional knockout of the homeobox genes *En1* and *En2* (*En1/2*) in the rhombic lip-derived eCN and granule cell precursors leads to embryonic loss of a subset of medial eCN and cell non-autonomous and location specific loss of Purkinje cells, with subsequent proportional scaling down of cortex growth. We propose that subsets of eCN dictate the survival of their specific Purkinje cell partners, and in turn sonic hedgehog secreted by Purkinje cells scales the expansion of granule cells and interneurons to produce functional local circuits and the proper folded morphology of the cerebellum.

## 1 Introduction

A key challenge for development of the brain is to produce the numerous cell types in the correct proportions from different stem cell pools at particular time points, and assembling the cells into nascent networks within brain regions that grow extensively during development. The cerebellum represents a powerful system in which to study this phenomenon since by birth the specification of neuronal lineages is complete but neurogenesis of several cell types has just begun. During the third trimester and first few months of human life the cerebellum undergoes a rapid expansion from a smooth ovoid anlage into a morphologically complex folded structure with specialized subregions. The folds (lobules) are considered the cerebellar cortex, and they overlay the cerebellar nuclei (CN), two bilaterally symmetrical groups of mediolaterally-arrayed nuclei in mice (medial, intermediate, and lateral nuclei), that house the main output neurons of the cerebellum. The layered cortex has an outer molecular layer, which contains interneurons and the axons of granule cells and dendrites of Purkinje cells (PCs), that is above a single layer of PC soma intermixed with the cell bodies of Bergmann glia. Below this layer is a dense layer of granule cells called the inner granule cell layer (IGL). PC axonal projections establish the only electrophysiological and physically direct connection between the cerebellar cortex and CN. The excitatory CN neurons (eCN) are the first to be born in the embryo, followed by the PCs and the interneurons that are their presynaptic partners. Postnatal cerebellar growth in the mouse is principally driven by the expansion of two progenitor populations that produce the presynaptic partners of PCs, granule cells and interneurons as well as astrocytes. Genes regulating differentiation of most cerebellar cell types have been identified, but the mechanisms responsible for coordinating the scaling of their cell numbers is poorly understood, despite their importance for understanding the formation of cerebellar circuit assembly.

During mouse cerebellar development, neurogenesis occurs in several germinal compartments: 1) the rhombic lip (RL) which produces the eCN between embryonic day (E) 9.5-E12.5 and then unipolar brush cells directly (Wang et al., 2005; Machold and Fishell, 2005; Sekerkova et al., 2004); 2) the ventricular zone which generates inhibitory PCs by E13.5 and early born interneurons including those that populate the CN (Sudarov et al., 2011; Leto and Rossi, 2012; Leto et al., 2016); 3) a RL-derived pool of granule cell precursors (GCPs) that migrates over the cerebellum and forms an external granule cell layer (EGL) from E15.5 to postnatal day (P) 15 that contains an outer layer of proliferating cells and inner layer of postmitotic granule cells; and 4) a ventricular zone-derived intermediate progenitor pool that expresses nestin and expands in the cerebellar cortex after birth and produces astrocytes, including specialized Bergmann glia, and late born interneurons of the molecular layer (Fleming et al., 2013; Parmigiani et al., 2015). The embryonic PCs form a multi-layer beneath the nascent EGL and project to the CN by E15.5 (Sillitoe et al., 2009). Once born, granule cells descend along Bergmann glial fibers to form the IGL, and synapse onto PC dendrites in the overlying molecular layer (Hatten and Heintz, 1994; Sillitoe and Joyner, 2007). PCs secrete sonic hedgehog (SHH) after E17.5, which drives the massive postnatal neurogenic phase by stimulating proliferation of GCPs and nestin-expressing progenitors (Corrales et al., 2004; Corrales et al., 2006; Lewis et al., 2004; Fleming et al., 2013; Parmigiani et al., 2015; De Luca et al., 2015; Wojcinski et al., 2017). PCs thus act in a dual role as a synaptic bridge between the two main RL-derived glutamatergic neuronal subtypes and as a developmental regulator of the cortex. Scaling of the proportions of postnatally born cells in the cerebellar cortex has been proposed to be regulated by the number of PCs and the amount of SHH they produce. However, how the number of CN neurons and PCs are proportioned properly during embryogenesis has not been addressed. Furthermore, it is not known whether PCs in the cerebellar cortex are dependent on CN neurons for their development or scaling.

The engrailed genes (*En1/2*) encoding homeobox transcription factors provide a powerful genetic entry point for studying developmental scaling of neuron types, since *En* mutants have a seemingly well-preserved cytoarchitecture despite suffering cerebellar hypoplasia that preferentially affects distinct lobules in each mutant (Millen et al., 1994; Sgaier et al., 2005; Cheng et al., 2010; Orvis et al., 2012). For example, specific loss of *En1/2* in the *Atohl*-expressing RL-lineage (*Atoh1-Cre; En1^lox/lox^; En2^lox/lox^* conditional knockouts, referred to as *Atoh-En1/2* CKOs) results in preferential loss of cerebellum volume in the vermis (medial cerebellum), with foliation defects restricted to the anterior and central vermis (Figure 1A; (Orvis et al., 2012)). As a basis for studying the roles of the *En1/2* genes in scaling of cerebellar neurons, we confirmed that the numbers of GCs, PCs, and molecular layer interneurons in the mutants are well scaled down relative to the decrease in cerebellar area. Despite the scaling of neurons in the cortex, we found that *Atoh-En1/2* CKOs have motor behavior defects. The primary defect in *Atoh-En1/2* CKOs was discovered to be loss of a subset of CN neurons after E15.5 that results in an ~50% decrease in eCN in the medial and intermediate nuclei. Our embryonic analysis showed that the early loss of eCN is accompanied by a cell non-autonomous reduction of PCs and a distruption of the eCN-to-PC ratio in *Atoh-En1/2* CKOs. Circuit mapping further revealed that the PCs in distinct lobules that are differentially depleted in the vermis of mutants project to different regions of the medial CN. We propose a model whereby the number of eCN neurons sets the growth potential of the cerebellar cortex through supporting survival of a balanced PC population that allows scaling of cortical neurons based on SHH expression.

**Figure 1.**
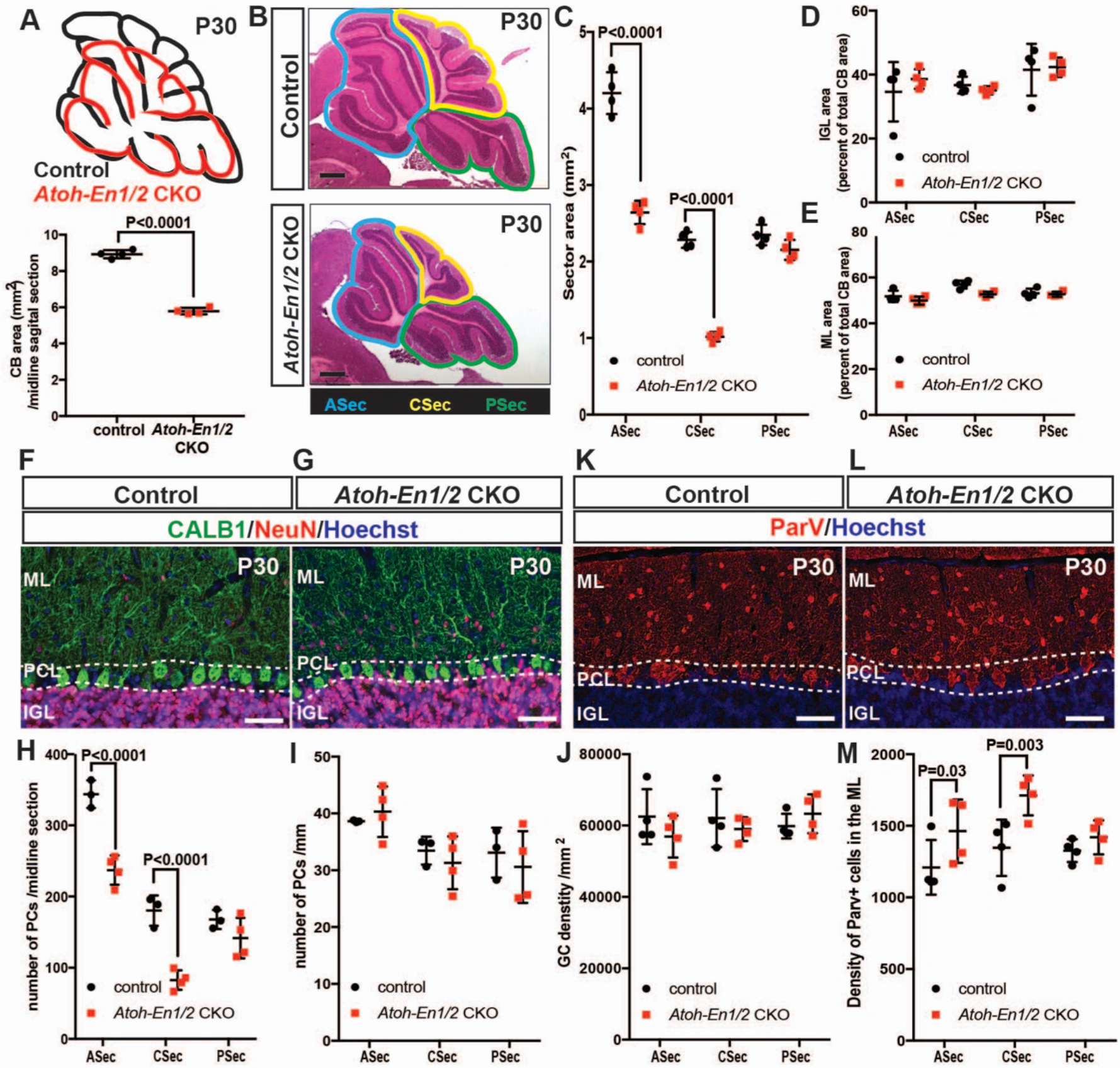
*En1/2* loss in the rhombic lineage results in preferential reduction of the anterior and central vermis while largely preserving scaling of cell types. **A)** Schematic representation (above) and quantification (below) of the average midline sagittal area of P30 *Atoh-En1/2* CKO and control cerebella (n=4 animals/condition, Student’s t-test). B) H&E staining of midsagittal sections of P30 mutant and control cerebella showing preferential reduction in the anterior (blue) and central (yellow) sectors (ASec and CSec) and not in the posterior (green)sector (PSec). **C)** Quantification of sector areas in P30 control and *Atoh-En1/2* CKO animals (n=4 animals/condition, Two-way ANOVA, F_(1,9)_=398.277, P<0.0001). **D-E)** IGL (D) and molecular layer (E) sector area quantifications as the percent of total average area showing no change in *Atoh-En1/2* CKOs compared to controls (n=4 animals/condition). **F-G)** Immunofluorescence analysis of P30 cerebellar sections for the PC marker calbindin1 (Calb1) and the pan-neuronal marker NeuN in an *Atoh-En1/2* CKO (G) compared to a control (F). **J-K)** Quantification of average PC numbers in each sector per midline sagittal section (H) showing reductions only in the ASec and CSec, whereas the density of PCs (K) is conserved (n=3 for controls and n=4 for *Atoh-En1/2* CKO, J: Two-way ANOVA, F_(115)_=72.52, P<0.0001). **L)** Quantification of granule cell density in each sector of mutants and controls (n=4 for each genotype). **K-L)** Immunofluorescence analysis of P30 cerebellar sections for the interneuron marker Parvalbumin (ParV) in an *Atoh-En1/2* CKO (K) compared to a control (L). **M)** Quantification of the density of ParV+ cells per sector in the molecular layer of mutants compared to controls (n=4 for each genotype, Two-way ANOVA, F_(19)_=28.4, P<0.0005). Significant *post hoc* comparisons are shown in the figure. CKO: conditional knockout, IGL: internal granule layer, ML: molecular layer, PCL: Purkinje cell layer, PC: Purkinje cell, GC: granule cell. Scale bars B: 500 μm, F-I: 100 μm.

## Results

### Loss of *En1/2* in the rhombic lip lineage results in preferential reduction of the anterior and central vermis but scaling of neurons in these regions is largely normal

Our previous study using 3D Magnetic Resonance Imaging showed a 25% reduction in the volume of the vermis of *Atoh-En1/2* CKOs but preserved cytoarchitecture (Orvis et al., 2012). In order to determine whether the *Atoh-En1/2* CKOs can serve as a useful model for studying scaling of neuron numbers during development, we first tested whether the proportions of different neurons in the areas of the vermis with reduced size were indeed normal. As a proxy for analyzing cerebellum size, we quantified the area of mid-sagittal sections of ~P30 animals, and found a 31.2 ± 2.0% reduction (n=4) in *Atoh-En1/2* CKOs compared to littermate controls (*En1^lox/lox^; En2^lox/lox^* mice) (Figure 1A). In order to determine whether particular regions of the vermis along the anterior-posterior axis were preferentially diminished in the mutants, we divided the cerebellum into three regions (Figure 1B) - an anterior sector (ASec), anterior to the primary fissure (lobules 1-5); a central sector (CSec), between the primary and secondary fissures (lobules 6-8); and a posterior sector (PSec), posterior of the secondary fissure (lobules 9-10) – and measured the areas of each. Interestingly, the CSec had the greatest decrease in area (55.4 ± 0.03%) with the ASec having a 37.1 ± 0.04% decrease and no significant reduction in the PSec of *Atoh-En1/2* CKOs compared to littermate controls (Figure 1C). Thus, *Atoh-En1/2* mutants have a differential reduction in the sizes of the central and anterior regions of the vermis.

We next determined whether the area of the layers compared to the total area of each sector and the numbers of each cell type within each sector were scaled proportionally in *Atoh-En1/2* CKOs. As predicted, the areas of the IGL and molecular layer were reduced only in the ASec and CSec and their proportions were conserved relative to the area of each sector in *Atoh-En1/2* CKOs (Figure 1D,E and Figure 1-figure supplement 1A-B). In addition, no change was observed in the areas of the IGL and molecular layer in the PSec of *Atoh-En1/2* CKOs. Furthermore, quantification of the number of PCs per midline section and PC density at ~P30 based on staining for Calbindin1, revealed that the PC numbers were reduced specifically in the ASec and Csec (31.0 ± 0.1% and 54.1 ± 0.1% reduction, respectively) but their densities were not altered in all sectors of *Atoh-En1/2* CKOs compared to controls (Figure 1F-K), indicating their numbers had scaled down properly. Similarly, quantification of NeuN-expressing granule cells showed that the densities of granule cells in the three sectors were not altered (Figure 1J). Furthermore, the ratio of PCs to granule cells was preserved in the three sectors of mutants compared to controls (Figure 1-figure supplement 1C). The density of Parvalbumin+ molecular layer interneurons was only mildly increased in the ASec and CSec of mutants (1.2 ± 0.2 and 1.3 ± 0.1 fold, respectively, Figure 1K-M) and the ratio of interneurons to PCs or granule cells was preserved in the three sectors of *Atoh-En1/2* CKOs compared to controls (Figure 1-figure supplement 1D-E). Thus, the major neuron types in midline sections of the cerebellar cortex are scaled down relative to the decrease in area and the cell-to-cell ratio of the cortical neurons of the CB is preserved. This result is consistent with the hypothesis that SHH secreted by each PC determines the the level of production of interneurons and granule cells that occupy the local territory. Indeed, quantification of the proportion of the EGL that contains the outer layer of proliferating GCPs at P6 showed that the ratio was preserved despite the EGL being smaller in the ASec and CSec (Figure 1-figure supplement 2).

### *Atoh1-En1/2* mutants have motor defecits

We next tested whether a smaller cerebellum that has a well scaled cortex can support normal motor behavior, since we had shown that irradiated mice with an ~20% reduction in the area of mid-sagittal sections but well scaled layers have no obvious motor defects (Wojcinski et al., 2017). Two motor behavior paradigms and grip strength were tested. Unlike irradiated mice, significant defects in motor behavior were observed in *Atoh-En1/2* CKOs. The performance of mutants on trials of the accelerating rotarod showed a significant difference on the third day, as well as the cumulative measurement of performance in *Atoh-En1/2* CKOs (160.1 ± 17.2 sec) compared to littermate controls (227.7 ± 18.4 sec) (Figure 2A). In the footprint analysis of gait, mutant mice had a significant decrease in stride length (6.3 ± 0.2 cm), a trend towards a decrease in stance length (4.1 ± 0.1 cm), and an increase in sway length (2.6 ± 0.1 cm) compared to controls (7.4 ± 0.2 cm stride; 4.5 ± 0.1 cm stance; 2.4 ± 0.1 cm sway)(Figure 2B-C). Curiously, *Atoh-En1/2* CKOs had a significant decrease in grip stength (normalized force= 4.9 ± 0.3) compared to littermate controls (normalized force= 6.3 ± 0.3) (Figure 2D). While the decrease in grip strength in mutants could contribute to the defects in motor behaviors, this is not likely the only factor as CNS-specific conditional mutants for *Cxcr4* have a decrease in grip strength but no change in stance (Huang et al., 2014) and mutants with greatly reduced grip strength can behave normally on the rotarod and have normal gait (Heck et al., 2008). Our finding that *Atoh1-En1/2* mutants have motor deficits could be because the cerebellum is even smaller than in the irradiated mice we studied, and/or that there is an additional defect outside the cerebellar cortex.

**Figure 2.**
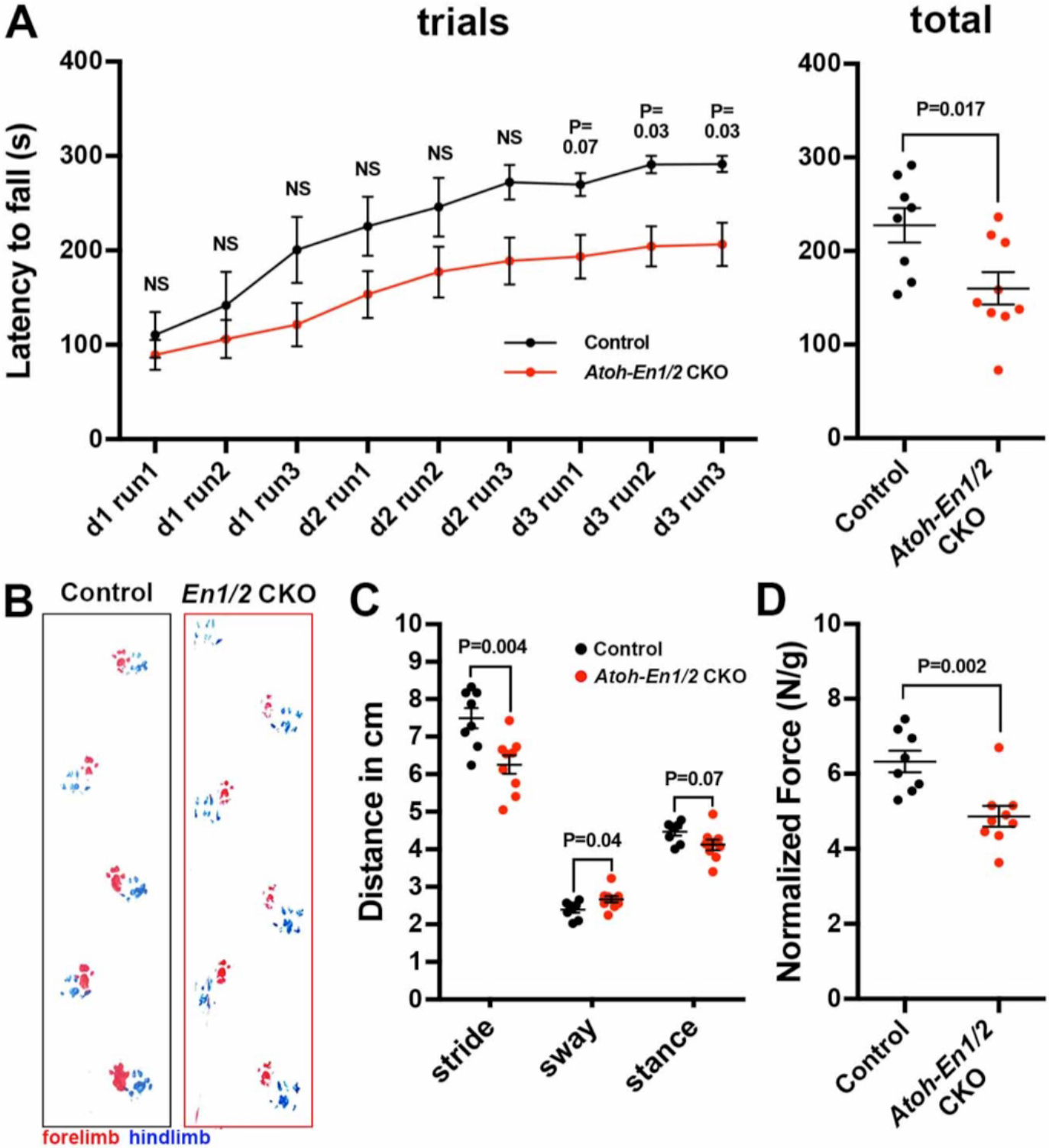
*Atoh-En1/2* CKO animals show motor behavior defects compared to control animals. **A)** Latency to fall from rotarod at each trial and total for control animals (n=8) compared to *Atoh-En1/2* CKOs (n=9). **B-C)** Representative images (B) and quantification (C) of footprint analysis performed on control (n=8) and *Atoh-En1/2* CKO (or *En1/2* CKO in B) (n=9) animals. **D)** Quantification of grip strength analysis of *Atoh-En1/2* CKO animals (n=9) compared to controls (n=8). Student’s t-test is shown in all panels.

### Loss of *En1/2* in the RL-lineage results in a preferential loss of medial and intermediate eCN neurons in the adult

Given that EN1/2 are expressed in the RL-derived eCN neurons that express MEIS2 (Wilson et al., 2011) and *Atoh-En1/2* mutants lack *En1/2* function in these neurons, a good candidate for an additional defect in the mutants was the CN projection neurons. In order to determine whether the loss *En1/2* in the RL-lineage results in any defect in the CN, we quantified the number of eCN in ~P30 mice. Stereology of every other section of coronal sections of half the cerebellum of *Atoh-En1/2* CKOs and littermate controls was used to quantify the number of large neurons in each nucleus based on Nissl staining, which include the eCN (Figure 3A-I). We confirmed that *Atoh-Cre* recombines in all the eCN by generating mice carrying *Atoh-Cre* and the *R26^LSL-nTDTom^* reporter allele (*Atoh-nTDTom* mice) and co-labeled for nuclear TDTom and three eCN markers, NeuN, and MEIS2 protein or *Slc17a6* RNA. All but rare TDTom+ cells expressed NeuN and the vast majority expressed *Slc17a6*, and very few cells expressed either marker and did not express TDTom (Figure 3-figure supplement 1A-O). Almost all TDTom+ cells also expressed MEIS2, although at intermediate levels along the medial-lateral axis the labeling was less extensive (Figure 3-figure supplement 1P-R). Furthermore, almost all cells expressing *Slc6a5* RNA, a marker for glycinergic cells, were negative for TDTom (Figure 3-figure supplement 1S).

**Figure 3.**
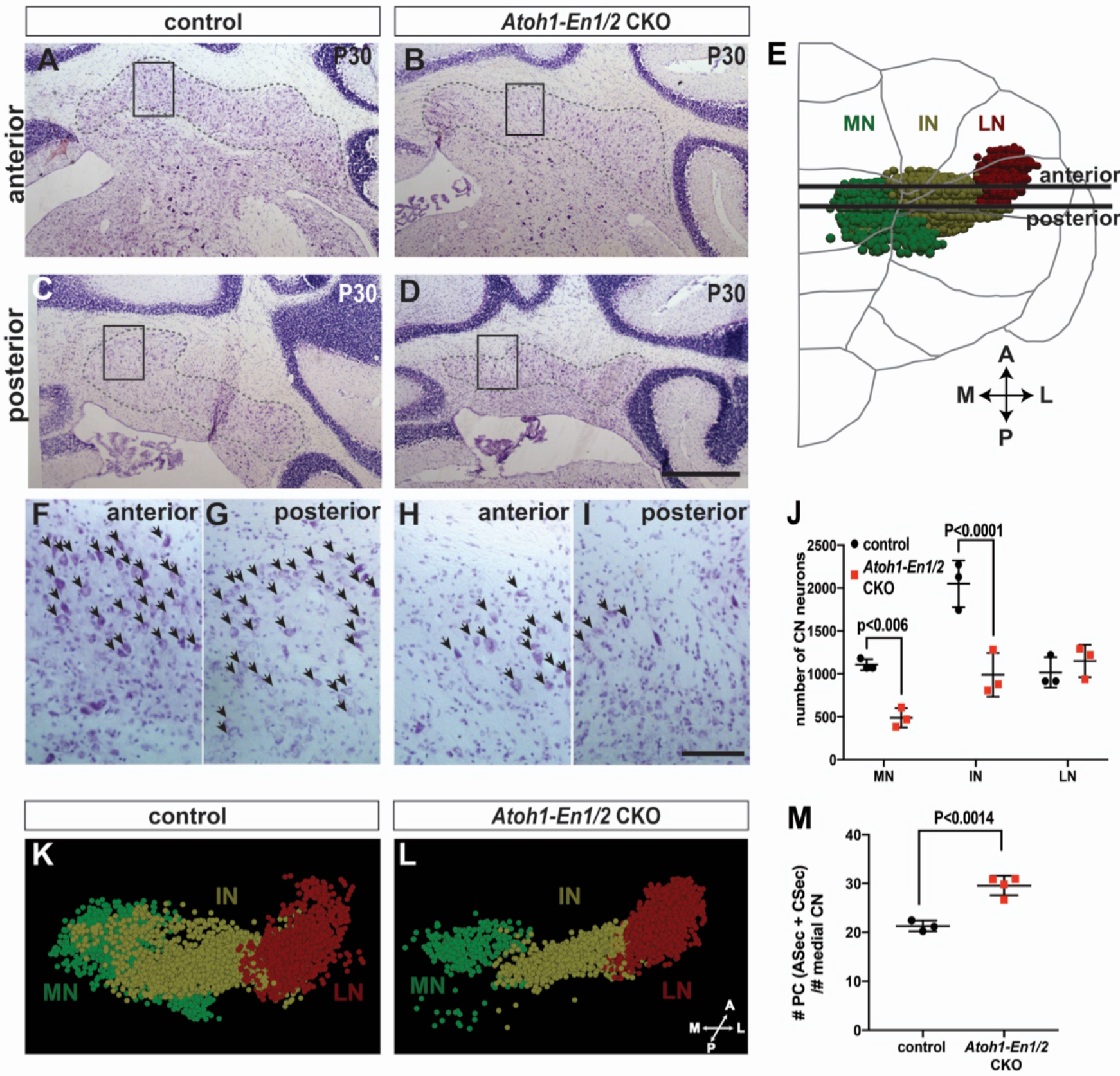
*Atoh-En1/2* CKO animals show preferential loss of medial and intermediate eCN and disrupted eCN-to-PC ratio. **A-D)** Nissl images of coronal sections from anterior (A-B) and posterior (C-D) cerebella of control (A and C) and *Atoh-EN1/2* CKO (B and D) animals with region of CN outlined by dotted lines. Boxes indicate higher magnification images shown in (FI). **E)** Schematic representation of a half brain with a 3D reconstruction of CN in a normal cerebellum. Stereology analysis was performed using similar coronal Nissl images. **F-I)** High magnification images of Nissl stained sections showing the loss of large eCN cells in *Atoh-En1/2* CKO animals (H and I) compared to controls (F and G). Arrows show the cells quantified for stereology analysis. **J)** Quantification of eCN neurons in the medial, intermediate, and lateral nuclei of the CN in one half of the *Atoh-En1/2* CKO cerebellum (n=3) compared to littermate controls (n=3) (Two-way ANOVA, F_(112)_=32.29, P=0.0001). Significant *post hoc* comparisons are shown in the figure. **K-L)** 3D reconstructions of stereology of the eCN in one half of the cerebellum from control (K) and *Atoh-En1/2* CKO (L) animals showing the reduction in medial and intermediate nuclei and a preferential loss of the posterior medial nucleus. **M)** Quantification of the ratio of PCs (from anterior and central sectors) to eCN in the medial nucleus as a measurement of PC-to-eCN neuron scaling in *Atoh-En1/2* CKO animals (n=4 for PCs and n=3 for eCN) compared to controls (n=3 for PCs and n=3 for eCN, Student’s t-test). MN: medial nucleus, IN: intermediate nucleus, LN: lateral nucleus. Scale bars A-D: 500 μm, F-I: 100 μm.

Interestingly, while there was an ~63% reduction in the total number of large neurons in the *Atoh-En1/2* CKOs compared to controls (n=3 each genotype), stereology revealed a marked loss of large neurons in the medial (control: 1107 ± 36.83; *Atoh-En1/2*: 487 ± 64.93) and intermediate nuclei (control: 2049 ± 157.4; *Atoh-En1/2:* 989 ± 147.1), whereas no alteration was detected in the lateral nuclei (control: 1017 ± 102.3; *Atoh-En1/2:* 1150 ± 108.5)(Figure 3J-L, Figure 3-supplement 2 and supplementary movies 1-2). Interestingly, within the medial CN, the posterior region appeared to be particularly diminished in *Atoh-En1/2* CKOs. We confirmed that the neurons lost in *Atoh1-En1/2* CKOs included eCN by generating a mutant carrying *Atoh1-Cre* and *R26^LSL-TDTom^* (*Atoh-TDTom-En1/2* CKO mutants) and stained sagittal cerebellar sections for NeuN and TDTom (Figure 3-figure supplement 2). NeuN+ cells were specifically reduced in the medial and intermediate CN, and many of the remaining NeuN+ cells were smaller in mutants than in controls. Thus, *En1/2* play a crucial role in the development of the two medial CN. Given that the two nuclei that are reduced in *Atoh-En1/2* CKOs specifically receive input from cortical structures that are also reduced in the mutants (vermis/paravermis lobules 1-8), this result suggests there could be a causal relationship between the loss of eCN and their PC presynaptic targets (Figure 1H).

As a means to assess whether scaling of PCs and eCN was preserved in adult mutants, we compared the decrease in the number of eCN in the medial nucleus of *Atoh-En1/2* CKOs to the number of PCs in the ASec and CSec per midline section, since the PCs in lobules 1-8 of the medial vermis project primarily to the medial nucleus. The PCs in much of lobule 9 and all of 10 project outside the cerebellum to the vestibular nuclei (Walberg and Dietrichs, 1988; Leto et al., 2016). Unlike the scaling of granule cells and interneurons to PCs in the cerebellar cortex, the number of medial eCN in *Atoh-En1/2* CKOs was reduced to 38% of that of controls, whereas the number of PCs in the midline was reduced to 55% (Figure 3M). The changes in neuron numbers resulted in an estimated 1.4 ± 0.1 fold increase in the ratio of the number of PCs to medial eCN in the ASec and CSec of *Atoh-En1/2* CKO animals compared to controls.

Our finding that in control mice there are ~20 PCs per eCN is in line with estimates that in rodents there is a functional convergenence of 20-50 PCs per eCN (Person and Raman, 2011). Thus, it appears that the PCs did not fully scale in proportion to the loss of medial eCN. Furthermore, the inbalance between the number of eCN and PCs might contribute to the motor defects in *Atoh1-En1/2* mutants.

### A cerebellum growth defect can be detected in *Atoh1-En1/2* CKO embryos

Given the possibility of a causal relationship between the reduction in the number of eCN and PCs in *Atoh-En1/2* adult mutants, we examined when a growth defect is first detected during cerebellum development, and whether the timing correlates with when eCN and/or PC numbers are reduced. Our previous analysis showed a clear growth and foliation defect in *Atoh-En1/2* mutants at P1 (Orvis et al., 2012); therefore, we analyzed embryonic stages (Figure 4A-C). Quantification of the cerebellar area of mid-sagittal sections at E17.5 of mutants and controls revealed a small reduction in area at E17.5 in *Atoh-En1/2* CKOs compared to littermate controls (13.3 ± 5.3 %) (Figure 4C). In addition, there appeared to be a delay in the formation of the earliest fissures at E17.5 (Figure 4A-B). Thus, a phenotype is first apparent in *Atoh-En1/2* CKOs when the main cell types that have been generated in the cerebellum are CN neurons and PCs, and long before the major expansion of GCPs.

**Figure 4.**
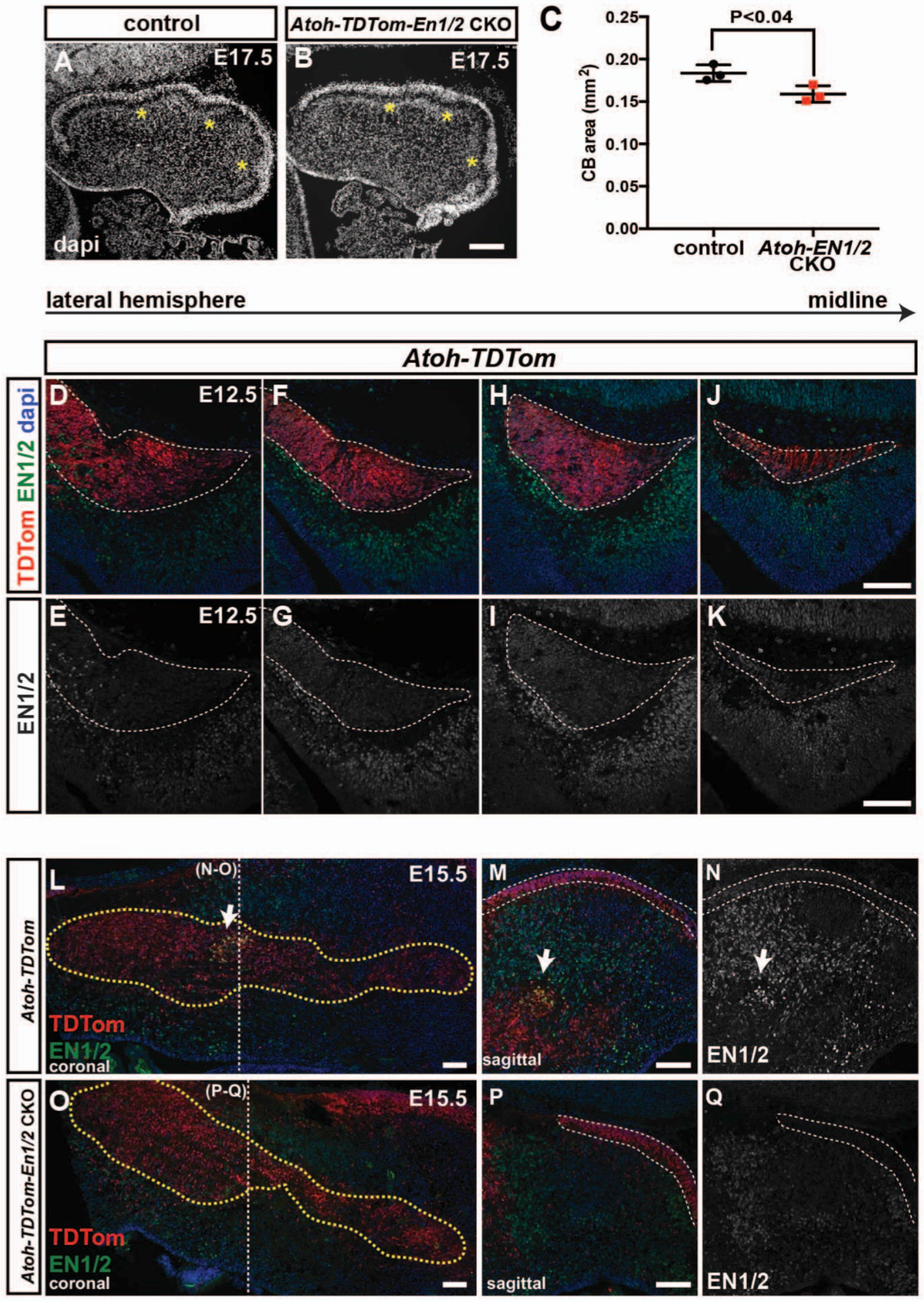
The cerebellum growth defect initiates embryonically when EN1/2 proteins are dynamically expressed in a subpopulation of eCN. **A-B)** Midsagittal sections of the cerebellum stained with the nuclear stain DAPI to show the morphology of the cerebellum at themidline of E17.5 *Atoh-En1/2* CKO animals (B) compared to controls (A). Asterisks indicate where the primary fissures are forming. **C)** Quantification of cerebellar area in sagittal sections at E17.5 showing a significant reduction in the cerebellar area of *Atoh-En1/2* CKO animals (n=3) compared to littermate controls (n=3, Student’s t-test). **D-K)** Lateral to medial sagittal sections showing the expression pattern of EN1/2 at E12.5 in *Atoh-TDTom* mice. Dotted white outlines show the nuclear transitory zone that is marked by TDTom. **L-Q)** EN1/2 expression in E15.5 *Atoh-TDTom* (L-N) and *Atoh-TDTom-EN1/2* CKO (O-Q) animals shows the loss of a subset of EN1/2-expressing CN cells (arrow). CN are outlined with yellow dotted lines. L,) Coronal sections. M-N and P-Q) Sagittal sections at the levels indicated by the dotted lines in (L, O). Loss of EN1/2 can also be observed in the EGL (outlined in white dotted lines). Scale bars: 100 μm.

### EN1/2 are dynamically expressed in the medial CN at E12.5-17.5

We next characterized the expression of EN1/2 proteins in the developing CN in more detail, as *En1/2* are expressed in subsets of all cerebellar cell types from as early as E8.5 (Davis et al., 1991; Millen et al., 1995; Wilson et al., 2011) and we previously only reported expression of the proteins in a small percentage of the medial MEIS2+ eCN at E17.5 (Wilson et al., 2011). An antibody that detects both EN1 and EN2 (Davis et al., 1991) was used to detect the two proteins in sagittal sections of the cerebellum in *Atoh-TDTom* mice at E12.5, 15.5, and 17.5 (Figure 4D-K and Figure 4-figure supplement 1). The specificity of the antibody was confirmed by showing absence of signal in the CN and EGL of *Atoh-En1/2* mutants (*Atoh-TDTom-En1/2* CKO mice) at E15.5 (Figure 4L-Q). We found that at E12.5, EN1/2 were weakly expressed in the TDTom+ cells of the forming EGL and nuclear transitory zone that houses the newly born CN, primarily in more medial sections (Figure 4J-K). In addition, EN1/2 were expressed in most of the PCs (Figure 4-figure supplement 1). By E15.5, EN1/2 expression was reduced in the EGL, CN and PCs, and in the CN was only detected in what is likely the precursors of the intermediate nucleus (Figure 4L-N).

### *En1/2* are required for survival of embryonic RL-derived eCN neruons

Given that a loss of eCN neurons could contribute to the phenotype of E17.5 *Atoh-En1/2* CKOs, we quantified the number of RL-derived eCN neurons that expressed the broad eCN marker MEIS2 and also TDTom in *Atoh-TDTom-En1/2* CKO and *Atoh-TDTom* controls (n=3 mice of each genotype) at E15.5 and E17.5. All MEIS2+ cells were labeled with cytoplasmic TDTom. In control mice, 67.7 ± 3.8% of the TDTom+ cells expressed MEIS2 at E15.5 and 63.5 ± 3.5% at E17.5, and a similar percentage of TDTom+ cells expressed MEIS2 in mutants (Figure 5-figure supplement 1A-B. None of the cells expressed the PC marker FOXP2 (Figure 5-figure supplement 1C-D). The TDTom+ cells in the EGL expressed PAX6, as well as a small number of cells in the forming IGL (Figure 5-figure supplement 1E-F). Quantification of the number of MEIS2+ or TDTom+ cells on every other sagittal section of half of the cerebellum revealed no significant change in the total number of MEIS2+ or TDTom+ cells at E15.5 in *Atoh-TDTom-En1/2* CKO compared to *Atoh-TDTom* controls (Figure 5A-B and Figure 5-figure supplement 2A-B). The distributions of the labeled cells along the medial-lateral axis were slightly altered in mutants compared to controls such that there was an ectopic population of eCN located near the midline and less cells laterally in mutants (Figure 5A,C-H and Figure 5-figure supplement 2A). At E17.5, the MESI2+ and TDTom+ cells in the CN of mutants were significantly reduced in number by 18.5 ± 1.6 % and 11.4 ± 2.1%, respectively (MEIS2+ mutant = 7957 ± 128.4 and control = 9766 ± 271.7; TDTOM+ mutant = 13649 ± 270.1, and control = 15407.3 ± 590.5, n=3/condition) (Figure 5I-J and Figure 5-figure supplement 2D-E). The distributions of the labeled cells along the medial-lateral axis showed a reduction in what will likely become the intermediate and medial nuclei (Figure 5I, K-P and Figure 5-figure supplement 2D). Given the loss of eCN between E15.5 and E17.5 in *Atoh1-En1/2* CKOs, we examined whether cell death was elevated in mutants compared to controls. TUNEL assay for cell death indeed indicated an increase in cell death in the region of the TDTom+ eCN population of E15.5 and E17.5 *Atoh-En1/2* mutants compared to controls (Figure 5-figure supplement 3). Thus, loss of *En1/2* in the RL-lineage results in a loss of eCN starting at ~E15.5.

**Figure 5.**
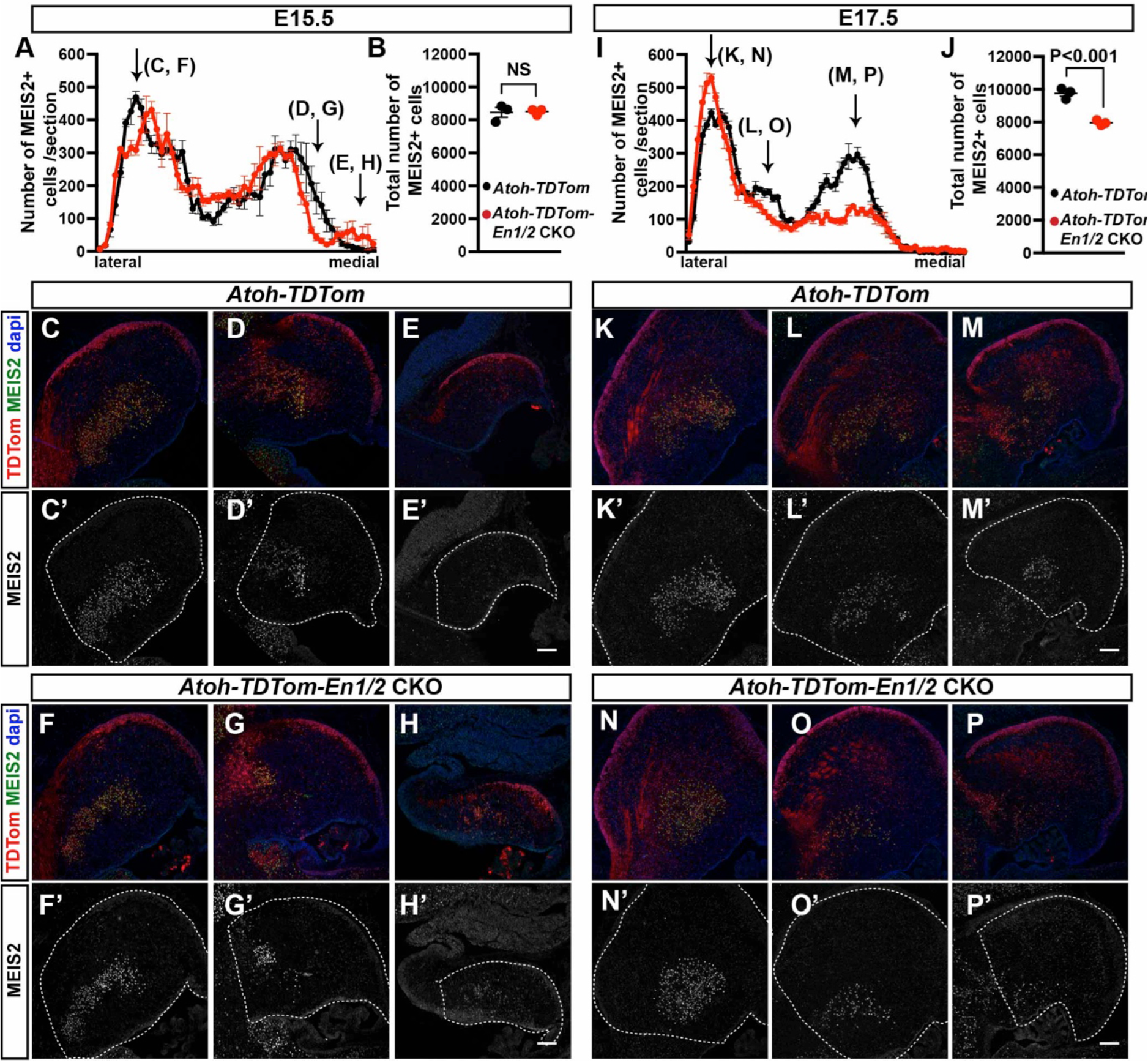
Loss of En1/2 leads to reduction of the medial RL-derived CN neurons. **A-B)** Quantification of the average number of MEIS2+ eCN cells in each sagittal section along the medial-lateral axis (A) and total eCN numbers (B) at E15.5 in *Atoh-TDTom* compared to *Atoh-TDTom-En1/2* CKOs (n=3 each genotype). **C-H)** Representative images of sagittal sections at the levels indicated by arrows in (A) stained for MEIS2 and TDTom in *Atoh-TDTom* (C-E) and *Atoh-TDTom-En1/2* CKOs (F-H). Although the total number of MEIS2+ cells is similar in mutants and controls (Student’s t-test, P=0.88), organization of the eCN is different in the mutant. **I-J)** Quantification of the average number of MEIS2+ eCN cells in each sagittal section along the medial-lateral axis (I) and total eCN numbers (J) at E17.5 in *Atoh-TDTom* compared to *Atoh-TDTom-En1/2* CKOs (n=3 for each genotype). **K-P)** Representative images of sagittal sections at the levels indicated by arrows in (I) stained for MEIS2 and TDTom in *Atoh-TDTom* (K-M) and *Atoh-TDTom-En1/2* CKOs (N-P). The total number of MEIS2+ cells (J) is reduced in the mutants compared to controls, specifically in the medial and intermediate sections (I, L, M) (n=3 for each genotype, Student’s t-test). Outlines show the regions of the cerebella. Scale bars: 100 μm.

### *En1/2* are required cell-nonoautonomously for survival of embryonic Purkinje cells

Given that eCN are lost between E15.5 and E17.5 in *Atoh-En1/2* mutants, we determined whether PCs are lost during the same time period. FOXP2 labeling of PCs revealed no decrease in the number of PCs per midline section or density of the cells at E15.5 in *Atoh-TDTom-En1/2* CKO mutants compared to *Atoh-TDTom* controls (Figure 6A-D). At E17.5, however, there was a significant decrease (18.3 ± 1.1%) in the number of PCs per section and density of the cells (Figure 6E-H). In summary, PC numbers in the vermis are reduced in parallel with the loss of medial/intermediate eCN between E15.5 and E17.5.

**Figure 6.**
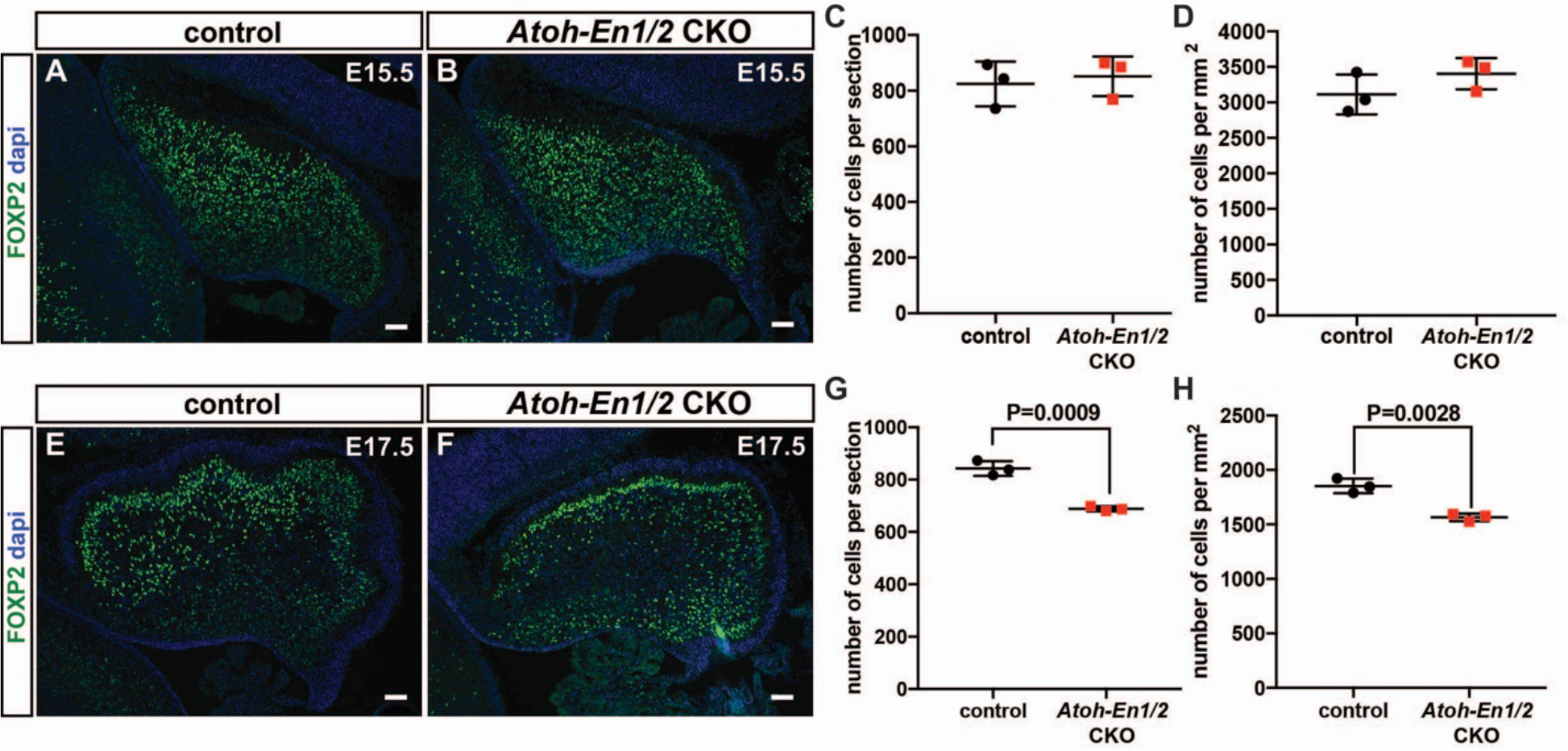
Purkinje cell loss occurs embryonically along with CN loss. **A-D)** Immunofluorescent analysis (A, B) of PCs (FoxP2+) and quantification of the number of PCs per midsagittal section (C) and their density (D) showing no significant changes between control and *Atoh-En1/2* CKO animals at E15.5 (n=3/condition). **E-H)** Immunofluorescent analysis (E, F) of PCs (FoxP2+) and the quantification of PCs (FoxP2+ cells) showing reduced PC numbers (G) and density (H) (n=3 each genotype) in *Atoh-En1/2* CKOs compared to controls at E17.5. Student’s t-test is shown in all panels.

### ASec and CSec PCs project to the distinct regions of the medial CN

We finally asked whether the PCs of the ASec and CSec preferentially project to distinct regions of the medial CN, since PCs broadly innervate the nuclei closest to them – medial vermis PCs project to the medial nuclei, hemispheric PCs project to the lateral nuclei, with the intermediate nucleus being innervated by the paravermis and lateral vermis (Leto et al., 2016). Although a comprehensive study has not been conducted as to whether PC axons project to the CN in a topographic manner based on their anterior-posterior position in lobules, neurons of the more posterior region of the medial CN, known as the fastigial oculomotor region, receive functional inputs from PCs in lobules 6 and 7 of the vermis (Zhang et al., 2016; Herzfeld et al., 2015; Person and Raman, 2011; Noda et al., 1990). As our data indicate that the posterior region of the medial CN is particularly diminished in *Atoh-En1/2* CKOs, and the size of the CSec is reduced to the greatest extent in these mutants, we asked whether axons from ASec and CSec PCs project to the anterior and posterior regions of the medial nucleus, respectively. Since most PCs of the PSec innervate the vestibular nucleus rather than the CN, we omitted the PSec from our CN-PC axon tracing analysis. We injected a Cre-inducible tracer virus (AAV5-EF1a-DIO-mCherry) into lobules 3-5, representing the ASec, or lobules 6/7 representing the CSec of ~P30 *Pcp2^Cre/+^* mice (Figure 7A-B), a mouse line that selectively expresses Cre recombinase in PCs (Zhang et al., 2004). 1.5 weeks post-infection, brains underwent delipidation to enhance immunohistology in the white matter (Renier et al., 2014) and 50 micron coronal sections were immunostained in a floating-stain method. Interestingly, we found that labeled PCs in the ASec preferentially innervated the anterior medial nucleus (Figure 7C-F) and labelled PCs from the CSec innervated the posterior medial nucleus (Figure 7C,G-I). The finding that ASec and CSec PCs preferentially innervate the anterior and posterior medial nucleus, respectively, parallels the regional differences in cerebellar hypoplasia seen in *Atoh1-En1/2* CKOs.

**Figure 7.**
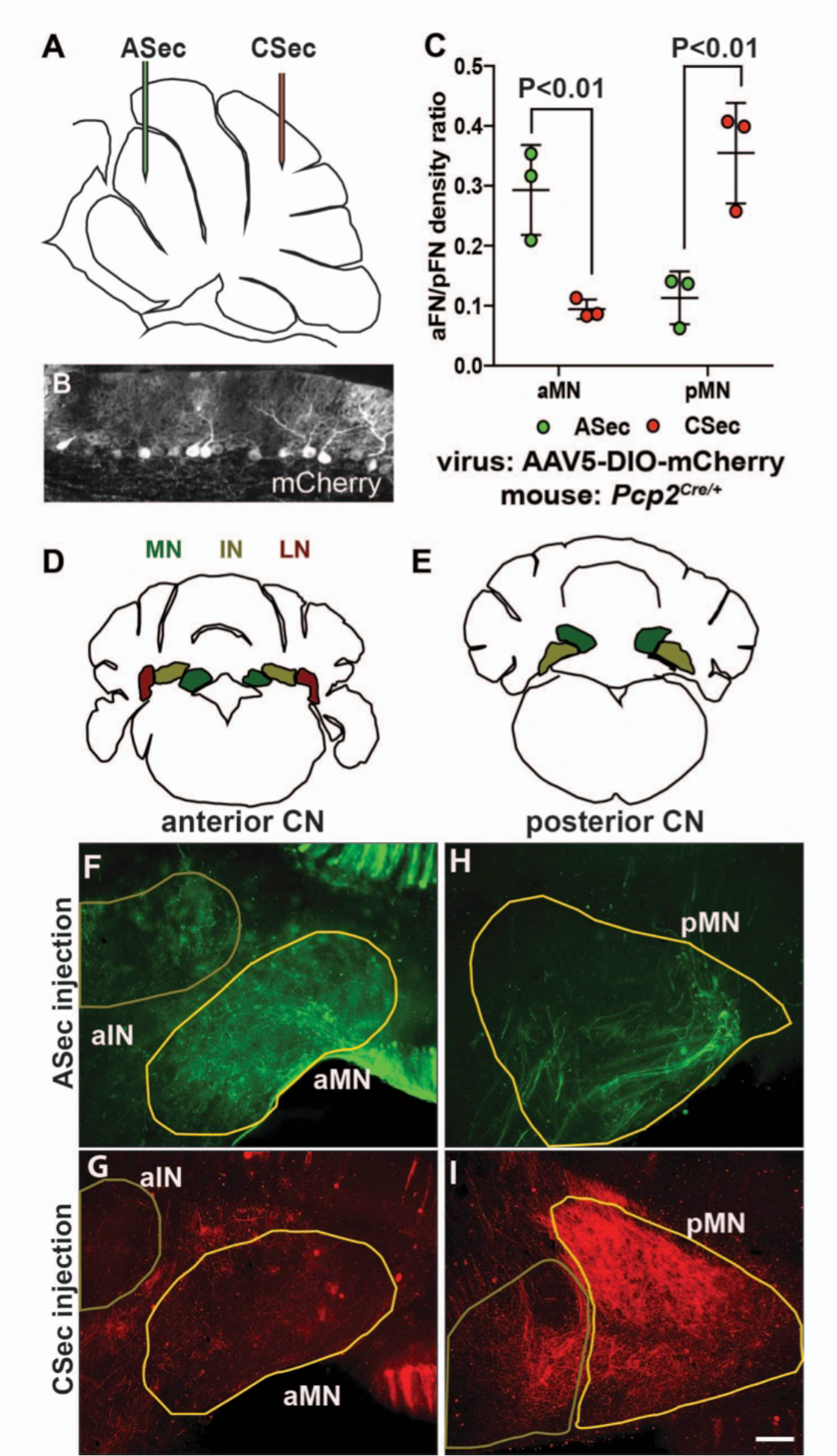
The anterior and central sectors of the vermis preferentially innervate different subregions of the medial CN. **A)** Diagram of scheme used for stereotactic injection of AAV-DIO-mCherry reporter virus to label PC axons projecting to the CN from the ASec or CSec of *Pcp2*^*Cre*/+^ animals. **B)** Sagittal section the PC layer of a *Pcp2*^*Crel*/+^ mouse injected with AAV-DIO-mCherry showing successful targeting of PCs. **C)** Quantification of fluorescence signal density in the anterior MN (aMN) and posterior MN (pMN) of ASec and CSec injected animals showing that ASec injection of virus preferentially innervates the aMN (n=3, Student’s t-test) and CSec labeled PCs preferentially innervate the pMN (n=3, Student’s t-test). **D-E)** Schematic representations of the anterior (D) and posterior (E) CN. **F-I)**. Immunofluorescent analysis showing the preferential labeling of the aMN or pMN in mice injected to trace axons projecting from PCs in the ASec (F-G) or CSec (H-I), respectively. Scale bar: 150 μm

## Discussion

In summary, we have uncovered that the cerebellum is constructed in a sequential order whereby the earliest-born neurons, the eCN, determine the cell number of their later-born pre-synaptic partners, PCs, likely based on secretion of a survival factor by eCN. In turn, the PC-derived mitogenic factor, SHH, then drives the production of all postnatally derived neurons and astrocytes of the cerebellar cortex (Figure 7-figure supplement 1) (Corrales et al., 2004; Corrales et al., 2006; Lewis et al., 2004; Fleming et al., 2013; Parmigiani et al., 2015; De Luca et al., 2015; Wojcinski et al., 2017). This progression of neurogenesis and maintenance of the overlying circuit constitutes a means by which developmental defects affecting the CN will lead to near scalar adaptation of cytoarchitecture in the cerebllar cortex. Nevertheless, we found that *Atoh1-En1/2* CKOs have motor behavior deficits, possibly because the PCs do not fully scale down to the number of eCN, and/or the smaller size of the cerebellum can not support robust long range neural circuits. Thus, both proper scaling and size regulation are necessary to generate robust neural circuits. The basic principles of our model can likely be applied to how circuits that span across brain regions could be scaled during development.

We found that scaling between the eCN and PCs does not appear to be as precise as the scaling within the cortex of *Atoh-En1/2* CKOs. One possibility for the ~1.4 fold higher ratio of PCs to eCN is that the difference reflects the mechanism by which scaling is attained. In the case of eCN:PC scaling, the process likely depends on the availability of a survival factor (such as a neurotrophin), whereas the PC:GC or PC:interneuron scaling depends on the amont of mitogen produced (SHH). An interesting additional possibility related to eCN:PC scaling is that since each PC projects to many eCN, if the survival of PCs is dependent on a factor secreted by eCN, then it might be that a greater proportion of PCs can survive than in normal homeostasis because some PCs continue to project to enough of the remaining eCN in mutants to receive sufficient survival factor. In contrast, the action of a mitogen is more likely to be concentration-dependent.

The results of our study indicate that if PCs are scaled back by birth by a specific genetic defect or by injury, then the other neurons generated postnatally in the cerebellar cortex will be scaled back in numbers to produce a normally proportioned cytoarchitecture. Considering the cerebellar cortex as a multilayer structure comprised of unitary tiles containing a single PC, which is able to support and specify postnatally-derived cells that contribute to its local circuit, the size of the PC pool can be thought of as a predictor of total postnatal cerebellum growth capacity. As the PCs redistribute into a monolayer sheet across the cerebellar cortex after birth, the smaller population of PCs in *Atoh-En1/2* CKOs is not able to spread out to form the normal length of the PC layer, since they have the same density at P30 as controls. The GCP pool is consequently reduced in size in mutants. As SHH expressed by PCs is required for expansion of the two proliferating precursor populations in the postnatal cerebellum, the scaling back in *Atoh-En1/2* CKOs of granule cells and interneurons, produced by GCPs and nestin-expressing progenitors, is likely due to the reduced amount of SHH present in the mutant cerebellum. In the context of the cerebellar cortex possibly being made up of repeated tiles of neurons, it is interesting to note that our results show that PCs regulate the numbers of their presynaptic neurons.

Based on our finding that PCs scale back in their numbers soon after many eCN die in late embryonic development in *Atoh-En1/2* CKOs, our study has provided evidence that the survival of PCs is dependent on CN. The timing of innervation of eCN by PCs fits with this proposal as axons from PCs extend into the CN by E15.5 (Sillitoe et al., 2009). Furthermore, *in vitro* assays have shown that PC survival is dependent on cell-cell interactions (Baptista et al., 1994) or on neurotrophins, and mouse mutants have shown that *Bdnf* and *p75^NTR^* null mutants exhibit cerebellar growth and foliation defects with some similarities to *Atoh1-En1/2* mutants (Schwartz et al., 1997; Carter et al., 2002; Carter et al., 2003). Based on gene expression in the Allen Institute Database and our analysis in *Atoh-TDTom* mice, *Bdnf* is expressed specifically in the CN at E15.5 (Figure 7-figure supplement 2); thus, it is a good candidate for a CN-derived survival factor for PCs. *Gdnf* is also detected in the CN after birth, suggesting that it could also contribute to long-term survival of PCs in the cerebellum.

As our model asserts that eCN neurons maintain the survival of their presynaptic PC neurons (Figure 7-figure supplement 1), we predicted that neurons in the medial nucleus that appear to be lost in the greatest numbers in the *Atoh-En1/2* CKOs should preferentially be innervated by PCs in the vermis lobules that also exhibit the most diminution. Stereotactic injection of AAV virus into the cerebellum to determine the projection pattern of labeled PC axons into the CN showed that PCs of the vermis central sector, where lobule growth and PC depletion are most pronounced, do indeed preferentially innervate the posterior region of the medial CN, which is preferentially reduced in cell number. In a complementary fashion, the PCs of the ASec preferentially project to the anterior medial nucleus and part of the intermediate nucleus.

Our study revealed that a primary defect in *Atoh-En1/2* CKOs is death of eCN starting at ~E15.5, indicating that *En1/2* are required in eCN for their survival in the embryo. *En1/2* have been shown to be required for survival of other neurons in the brain (Fox and Deneris, 2012; Sonnier et al., 2007). Similar to ablation of *En1/2* in eCN as they are being generated in the embryo, conditional knockout of *En1/2* in embryonic postmitotic serotonergic (5-HT) neuron precursors of the dorsal Raphe nucleus (DRN) results in a normal number of 5-HT neurons being initially generated, but the cells then become disorganized through abnormal migration pathways and undergo significant cell death around birth (Fox and Deneris, 2012). Our study of the eCN in *Atoh-En1/2* CKOs indicates a similar role for EN proteins in the settling of eCN into three nuclei and in their survival.

*En1/2* are ablated not only in the eCN of the cerebellum of *Atoh-En1/2* mutants, but also in the GCPs, which undergo their major expansion after birth. Some of the postnatal growth defects could therefore be due to a minor requirement for *En1/2* after birth in the GCPs, although we found that the EGL scales back in size but preserves a normal ratio of proliferating GCPs in the EGL. We nevertheless did recently show that deletion of *En1/2* in GCPs along with over-activation of SHH signaling results in a small increase in mutant cells remaining in the proliferative GCP pool, suggesting *En1/2* could play a minor role in promoting differentiation of GCPs (Tan et al., 2018). In addition to the RL-lineage, *Atoh1-Cre* deletes *En1/2* in the precerebellar nuclei, some of which express *En1/2.* Thus, some of the phenotype could be a secondary (cell-nonautonomous) effect of loss of *En1/2* in structures outside the cerebellum. If this is the case, it will be interesting to determine whether the loss of eCN is due in part to lack of a survival factor that is normally secreted by presynatic partners.

In conclusion, *En1* and *En2* are required soon after the eCN are produced by the RL for their survival, and possibly for proper settling into three bilaterally symmetrical CN. Loss of a subset of medial/intermediate eCN neurons results in cell non-autonomous loss of their specific presynaptic PC partners. The size of the cerebellar cortex is then scaled back to be proportional to the number of PCs that remain in each region of the vermis, producing a normal appearing cytoarchitecture. The PCs do not scale back fully, perhaps accounting for the motor behavior defects in *Atoh-En1/2* mutants. Our results have important implications for how scaling of neurons in all brain regions could be regulated by a combination of survival factors and mitogens, as well as for synaptic partners of neurons located in distant regions of the nervous system.

## Materials and methods

### Mouse strains

All animal experiments were performed in accordance with the protocols approved and guidelines provided by the Memorial Sloan Kettering Cancer Center’s Institutional Animal Care and Use Committee (IACUC). Animals were given access to food and water *ad libitum* and were housed on a 12-hour light/dark cycle.

All mouse lines were maintained on a Swiss-Webster background: *En1^lox^* (Sgaier et al., 2007), *En2^lox^* (Cheng et al., 2010), *Atoh1-Cre* (Matei et al., 2005), *R26^LSL-TDTom^* (ai14, Jackson labs stock no: 007909)(Madisen et al., 2010), *R26^LSL-nTDTom^* (ai75D, Jackson labs stock no: 025106), *Pcp2^Cre^* (Zhang et al., 2004). Noon of the day that a vaginal plug was discovered was designated developmental stage E0.5. Both sexes were used for all experiments and no randomization was used. Exclusion criteria for data points were sickness or death of animals during the testing period. For behavioral testing, investigators were blinded to genotype during the data collection and analyses.

### Behavioral Testing

Five-week-old animals (Control: n=8 and *Atoh-En1/2:* n=9) were used to for all of the motor behaviour testing described below.

*Rotarod:* Analysis was performed 3 times a day on 3 consecutive days. Animals were put on an accelerating rotarod (47650, Ugo Basile), and were allowed to run till the spead reached 5 rpm. Then, the rod was accelerated from 5-40 rpm over the couse of 300 seconds. Latency to fall was recorded as the time of falling for each animal. Animals rested for 10 minutes in their home cage between each trial.

*Grip Strength:* A force gauge with a horizontal grip bar (1027SM Grip Strength meter with single sensor, Columbus Instruments) was used to measure grip strength. Animals were allowed to hold the grip bar while being gently pulled away by the base of their tail. Five measurements with 5 minute resting intervals were performed for each animal and the data was reported as the average of all trials, normalized to the mouse’s weight (Force/gram).

*Footprinting Analysis:* After painting the forefeet and hindfeet with red and blue nontoxic acrylic paint (Crayola), respectively, animals were allowed to walk on a strip of paper. Experiments were performed in a 50 cm long and 10 cm wide custom-made Plexiglass tunnel with a dark box at one end. Each mouse was run through the tunnel 3 times and the distance between the markings were measured and averaged.

### Genotyping

The DNA primers used for genotyping each allele and transgene were as follows: *En1^lox^* (Orvis et al., 2012)), *En2^lox^* (Orvis et al., 2012), Cre (*Atoh1-Cre* and *Pcp2^Cre^*): 5’-TAAAGATATCTCACGTACTGACGGTG, 5’-TCTCTGACCAGAGTCATCCTTAGC; *R26^tdTom^* (nuclear and cytoplasmic): 5’-CTGTTCCTGTACGGCATGG, 5’-GGCATTAAAGCAGCGTATCC.

### Tissue Preparation and Histology

The brains of all embryonic stages were dissected in cold PBS and immersion fixed in 4% PFA for 24 hours at 4 °C. All postnatal stages were transcardial perfused with PBS followed by 4% PFA, and brains were post-fixed in 4% PFA overnight, and washed 3 times in PBS. Specimens for paraffin embedding were put through an ethanol-xylene-paraffin series, then embedded in paraffin blocks, and sectioned to 10 μm on a microtome (Leica Instruments). Specimens for cryosectioning were placed in 30% sucrose/PBS until they sank, embedded in OCT (Tissue-Tek), frozen in dry ice cooled isopentane, and sectioned at 14 μm on a cryostat unless otherwise stated (Leica, CM3050S).

### Immunohistochemistry

Sections of cryosectioned tissues were immersed in PBS for 10 min. After these initial stages of processing, specimens were blocked with blocking buffer (5% Bovine Serum Albumin (BSA, Sigma) in PBS with 0.2% Triton X-100). Primary antibodies in blocking buffer were placed on slides overnight at 4 °C, washed in PBS with 0.2% Triton X-100 (PBST) and applied with secondary antibodies (1:1000 Alexa Fluor-conjugated secondary antibodies in blocking buffer) for 1 hour at room temperature. Counterstaining was performed using Hoechst 33258 (Invitrogen). The slides were then washed in PBST, then mounted with a coverslip and Fluorogel mounting medium (Electron Microscopy Sciences).

Haematoxylene and Eosin (H&E) staining was performed for histological analysis and cerebellar area measurements, except at E17.5 where DAPI staining was used.

Nissl staining was used for sterology analysis in the eCN in adult brains.

### Antibodies

The following antibodies were used at the listed concentrations: Rabbit anti-MEIS2 (1:3000 for embryonic tissue, 1:2000 for adult sections, K846 rabbit polyclonal; (Mercader et al., 2005), Goat anti-FOXP2 (Everest Cat no: EB05226, 1:1000), Rabbit anti-panEN; 1:200 (Davis et al., 1991), anti-Calbindin (Swant Inc. Cat No: CB38, 1:1000 rabbit, Cat No: 300, 1:500 mouse), Mouse α-NEUN (Millipore Cat no: MAB377, 1:1000), mouse α-Parvalbumin (Millipore Cat no: MAB1572, 1:500)

### RNA *in situ* hybridization

Probes were *in vitro* transcribed from PCR-amplified templates prepared from cDNA synthesized from postnatal whole brain or cerebellum lysate. The primers used for PCR amplification were: *Slc17a6:* Forward: 5’-AGACCAAATCTTACGGTGCTACCTC-3’, Reverse: 5’-AAGAGTAGCCATCTTTCCTGTTCCACT-3’ *Slc6a5:* Forward: 5’-GTATCCCACGAGATGGATTGTT-3’, Reverse: 5’-CCATACAGAACACCCTCACTCA-3’ *Bdnf*: Forward: 5’-CGACGACATCACTGGCTG-3’, Reverse: 5’-CGGCAACAAACCACAACA-3’.

Primers were flanked in the 5’ with SP6 (antisense) and T7 (sense) promoters. Specimen treatment and hybridization were performed as described previously (Blaess et al., 2011).

### Image Acquisition and Analysis

All images were collected with a DM6000 Leica fluorescent microscope or a Zeiss Axiovert 200 with Apotome and processed using ImageJ (NIH) or Photoshop (Adobe). Image quantification was performed with ImageJ.

For the cerebellar, sector, IGL, and ML area, H&E stained slides were used. A region of interest was defined by outlining the perimeter of the outer edges of the region quantified. Three midline parasagittal sections/brain were analyzed and values were averaged for each brain.

Cell counts were obtained using the Cell Counter plugin for ImageJ (NIH). PC numbers were averaged from 3 midline parasagittal sections/brain. PC density was calculated by dividing the number of PCs by the length of the PCL. Granule cell density was measured by counting the NeuN+ cells in a 40x frame of the IGL from 3 midline parasagittal sections/brain and dividing the number of cells by the area counted. GC density for different sectors was obtained from lobules 3 (anterior), 6-7 (central), and 9 (posterior). GC numbers/midline parasagittal sections were calculated by multiplying the GC density by the area of the IGL. Parvalbumin+ cell counts in the ML were obtained from lobules 3 (anterior), 6-7 (central), and 9 (posterior) at the midline and the number of cells were divided by the ML area measured. Due to the high variability of the cell counts between sections, 5 midline parasagittal sections were quantified for this purpose. The number of ML Parvalbumin+ interneurons was extrapolated by multiplying the density with the ML area at the midline parasagittal sections.

Ratios of the different CB neurons with respect to each other were calculated using the numbers obtained for parasagittal midline sections as described above. The ratio of PCs to eCN was obtained by dividing the number of PCs determined above on midline sagital sections/brain from control and mutant animals at P30 by the average number of eCN at P30 from a different cohort of control and mutant mice.

### Stereology

Brains were paraffin embedded, as described above, and sectioned coronally at 10 μm. The sections were Nissl stained and every other section was analyzed to prevent double counting neurons split in serial sections. For each analyzed section in sequential order from rostral to caudal, the section perimeter of the CN on one half of the cerebellum was traced and each large projection neuron was registered. Sections were then aligned into a 3D representation in NeuroLucida.

### Stereotactic injection

One-month-old mice were anesthetized by isoflurane inhalation and head fixed in a stereotactic frame (David Kopf Instruments) with an isoflurane inhaler. The head of the animal was shaved, disinfected with ethanol and betadine, and a midline incision was made from between the eyes to the back of the skull. Coordinate space for the system was calibrated by recording the coordinates of Bregma and Lambda. After a small hole was drilled in the skull over the injection site, the needle was robotically injected into the cerebellum position specified by atlas coordinate (Neurostar StereoDrive). For targeting the ASec and PSec, coordinates of the injection were −6.00mm from bregma, −2.2mm deep from dura, and 8.5mm from bregma, 4.2mm deep from dura, respectively. One μL of 10^12^Tu/mL AAV5 virion in PBS was injected in 20 millisecond pulses at 10 psi with a Picospritzer. The needle was left *in situ* for 5 minutes, and then removed by 50 μm increments over about a minute. The scalp was then sealed by Vetbond adhesive, 0.1 μg/g of buprenorphine was administered for postoperative analgesia, and the animal was placed in a heated chamber for recovery. Brains from these animals were analyzed 1.5 weeks after surgery. Brains were processed for cryosectioning as above but were sectioned at 30 μm. To establish the accuracy of the injections to the appropriate sectors, *R26^tdTom/tdTom^* P30 animals were injected stereotactically with either AAV5-pgk-Cre virus or trypan blue dye in the midline of either the ASec (lobules 3-5) or CSec (lobules 6-7). Whole mount imaging of TDTom signal verified that the transduced cells remained restricted to the injection area and spread minimally in the lateral axis beyond the vermis. The tracing was done by injecting AAV5-EF1a-DIO-mCherry virus into *Pcp2^Crel+^* animals. The AAV-EF1a-DIO-mCherry plasmid was constructed by Karl Deisseroth and virus was packaged as serotype 5 and prepared to 10^12^ Tu/mL by the UNC Chapel Hill Vector Core.

### Tissue Delipidation for Tracing Analysis

After perfusion, postfix, and PBS washes as described above, brains were delipidated with a modified Adipo-Clear procedure (Chi et al., 2018) to enhance immunolabeling in heavily myelinated regions. The brain samples were washed with a methanol gradient (20%, 40%, 60%, 80%) made by diluting methanol with B1n buffer (0.1% Triton X-100/0.3 M glycine in H2O, pH 7.0), 30 min each step; then washed in 100% methanol for 30 min twice, and reverse methanol/B1n gradient (80%, 60%, 40%, 20%) for 30 min each; then in B1n buffer for 30 min twice. After delipidation, samples were washed in PBS and sectioned with a vibratome at 50 μm. The free-floating sections were immunolabeled for GFP and mCherry for analysis of PC projections.

### Statistical Analysis

Prism (GraphPad) was used for all statistical analyses. Statistical comparisons used in this study were Student’s two-tailed t-test and Two-way analysis of variance (ANOVA), followed by post hoc analysis with Tukey’s test for multiple comparisons. Relevant F-statistics and p-values are stated in the figure legends and the p-values of the relevant *post hoc* multiple comparisons are shown in the figures. A summary of all of the statistical analyses performed can be found in Figure 1_source data 1. The statistical significance cutoff was set at P<0.05. Population statistics were represented as mean ± standard deviation (SD) of the mean, except for the behavioural data presented in Figure 2. Population statistics in Figure 2 are represented as mean ± standard error (SE). No statistical methods were used to predetermine the sample size, but our sample sizes are similar to those generally employed in the field. n≥3 mice were used for each experiment and the numbers for animals used for each experiment are stated in the figure legends.

## Acknowledgements

We thank past and present members of the Joyner laboratory for discussions and technical help. We are grateful to Dr. Miguel Torres for providing a polyclonal antibody to all isoforms of mouse MEIS2. This work was supported by grants from the NIH to ALJ (R37MH085726) and RW (5F32NS080422), and a National Cancer Institute Cancer Center Support Grant [P30 CA008748-48]. Z.W. was supported by the Kavli Neural Systems Institute at the Rockefeller University.

## References

Baptista, C. A., Hatten, M. E., Blazeski, R. & Mason, C. A. 1994. Cell cell interactions influence survival and differentiation of purified Purkinje cells in vitro. Neuron, 12, 243–260.

Blaess, S., Bodea, G. O., Kabanova, A., Chanet, S., Mugniery, E., Derouiche, A., Stephen, D. & Joyner, A. L. 2011. Temporal-spatial changes in Sonic Hedgehog expression and signaling reveal different potentials of ventral mesencephalic progenitors to populate distinct ventral midbrain nuclei. Neural Dev, 6, 29. 1749-8104-6-29 [pii] 10.1186/1749-8104-6-29 [doi]

Carter, A. R., Berry, E. M. & Segal, R. A. 2003. Regional expression of p75NTR contributes to neurotrophin regulation of cerebellar patterning. Mol Cell Neurosci, 22, 113.

Carter, A. R., Chen, C., Schwartz, P. M. & Segal, R. A. 2002. Brain-derived neurotrophic factor modulates cerebellar plasticity and synaptic ultrastructure. J Neurosci, 22, 1316–27.

Cheng, Y., Sudarov, A., Szulc, K. U., Sgaier, S. K., Stephen, D., Turnbull, D. H. & Joyner, A. L. 2010. The Engrailed homeobox genes determine the different foliation patterns in the vermis and hemispheres of the mammalian cerebellum. Development, 137, 519–29. 10.1242/dev.027045

Chi, J., Wu, Z., Choi, C. H. J., Nguyen, L., Tegegne, S., Ackerman, S. E., Crane, A., Marchildon, F., Tessier-Lavigne, M. & Cohen, P. 2018. Three-Dimensional Adipose Tissue Imaging Reveals Regional Variation in Beige Fat Biogenesis and PRDM16-Dependent Sympathetic Neurite Density. Cell Metab, 27, 226–236.e3. 10.1016/j.cmet.2017.12.011

Corrales, J. D., Blaess, S., Mahoney, E. M. & Joyner, A. L. 2006. The level of sonic hedgehog signaling regulates the complexity of cerebellar foliation. Development, 133, 1811–21. 10.1242/dev.02351

Corrales, J. D., Rocco, G. L., Blaess, S., Guo, Q. & Joyner, A. L. 2004. Spatial pattern of sonic hedgehog signaling through Gli genes during cerebellum development. Development, 131, 5581–90. 10.1242/dev.01438

Davis, C. A., Holmyard, D. P., Millen, K. J. & Joyner, A. L. 1991. Examining pattern formation in mouse, chicken and frog embryos with an En-specific antiserum. Development, 111, 287–98.

De Luca, A., Parmigiani, E., Tosatto, G., Martire, S., Hoshino, M., Buffo, A., Leto, K. & Rossi, F. 2015. Exogenous sonic hedgehog modulates the pool of GABAergic interneurons during cerebellar development. Cerebellum, 14, 72–85. 10.1007/s12311-014-0596-x

Fleming, J. T., He, W., Hao, C., Ketova, T., Pan, F. C., Wright, C. C., Litingtung, Y. & Chiang, C. 2013. The Purkinje neuron acts as a central regulator of spatially and functionally distinct cerebellar precursors. Dev Cell, 27, 278–92. 10.1016/j.devcel.2013.10.008

Fox, S. R. & Deneris, E. S. 2012. Engrailed is required in maturing serotonin neurons to regulate the cytoarchitecture and survival of the dorsal raphe nucleus. J Neurosci, 32, 7832–42. 10.1523/jneurosci.5829-11.2012

Hatten, M. E. & Heintz, N. 1994. Mechanisms of Neural Patterning and Specification in the Developing Cerebellum. Annual Reviews.

Heck, D. H., Zhao, Y., Roy, S., Ledoux, M. S. & Reiter, L. T. 2008. Analysis of cerebellar function in Ube3a-deficient mice reveals novel genotype-specific behaviors. Hum Mol Genet, 17, 2181–9. 10.1093/hmg/ddn117

Herzfeld, D. J., Kojima, Y., Soetedjo, R. & Shadmehr, R. 2015. Encoding of action bythe Purkinje cells of the cerebellum. Nature, 526, 439–42. 10.1038/nature15693

Huang, G. J., Edwards, A., Tsai, C. Y., Lee, Y. S., Peng, L., Era, T., Hirabayashi, Y., Tsai, C. Y., Nishikawa, S., Iwakura, Y., Chen, S. J. & Flint, J. 2014. Ectopic cerebellar cell migration causes maldevelopment of Purkinje cells and abnormal motor behaviour in Cxcr4 null mice. PLoS One, 9, e86471. 10.1371/journal.pone.0086471

Leto, K., Arancillo, M., Becker, E. B., Buffo, A., Chiang, C., Ding, B., Dobyns, W. B., Dusart, I., Haldipur, P., Hatten, M. E., Hoshino, M., Joyner, A. L., Kano, M., Kilpatrick, D. L., Koibuchi, N., Marino, S., Martinez, S., Millen, K. J., Millner, T. O., Miyata, T., Parmigiani, E., Schilling, K., Sekerkova, G., Sillitoe, R. V., Sotelo, C., Uesaka, N., Wefers, A., Wingate, R. J. & Hawkes, R. 2016. Consensus Paper: Cerebellar Development. Cerebellum, 15, 789–828. 10.1007/s12311-015-0724-2

Leto, K. & Rossi, F. 2012. Specification and differentiation of cerebellar GABAergic neurons. Cerebellum, 11, 434–5. 10.1007/s12311-011-0324-8

Lewis, P. M., Gritli-Linde, A., Smeyne, R., Kottmann, A. & Mcmahon, A. P. 2004. Sonic hedgehog signaling is required for expansion of granule neuron precursors and patterning of the mouse cerebellum. Dev Biol, 270, 393–410. 10.1016/j.ydbio.2004.03.007

Machold, R. & Fishell, G. 2005. Math1 is expressed in temporally discrete pools of cerebellar rhombic-lip neural progenitors. Neuron, 48, 17–24. 10.1016/j.neuron.2005.08.028

Madisen, L., Zwingman, T. A., Sunkin, S. M., Oh, S. W., Zariwala, H. A., Gu, H., Ng, L. L., Palmiter, R. D., Hawrylycz, M. J., Jones, A. R., Lein, E. S. & Zeng, H. 2010. A robust and high-throughput Cre reporting and characterization system for the whole mouse brain. Nat Neurosci, 13, 133–40. 10.1038/nn.2467

Matei, V., Pauley, S., Kaing, S., Rowitch, D., Beisel, K. W., Morris, K., Feng, F., Jones, K., Lee, J. & Fritzsch, B. 2005. Smaller inner ear sensory epithelia in Neurog 1 null mice are related to earlier hair cell cycle exit. Dev Dyn, 234, 633–50. 10.1002/dvdy.20551

Mercader, N., Tanaka, E. M. & Torres, M. 2005. Proximodistal identity during vertebrate limb regeneration is regulated by Meis homeodomain proteins. Development, 132, 4131-42. 10.1242/dev.01976

Millen, K. J., Hui, C. C. & Joyner, A. L. 1995. A role for En-2 and other murine homologues of Drosophila segment polarity genes in regulating positional information in the developing cerebellum. Development, 121, 3935–45.

Millen, K. J., Wurst, W., Herrup, K. & Joyner, A. L. 1994. Abnormal embryonic cerebellar development and patterning of postnatal foliation in two mouse Engrailed-2 mutants. Development, 120, 695–706.

Noda, H., Sugita, S. & Ikeda, Y. 1990. Afferent and efferent connections of the oculomotor region of the fastigial nucleus in the macaque monkey. J Comp Neurol, 302, 330–48. 10.1002/cne.903020211

Orvis, G. D., Hartzell, A. L., Smith, J. B., Barraza, L. H., Wilson, S. L., Szulc, K. U., Turnbull, D. H. & Joyner, A. L. 2012. The engrailed homeobox genes are required in multiple cell lineages to coordinate sequential formation of fissures and growth of the cerebellum. Dev Biol, 367, 25–39. 10.1016/j.ydbio.2012.04.018

Parmigiani, E., Leto, K., Rolando, C., Figueres-Onate, M., Lopez-Mascaraque, L., Buffo, A. & Rossi, F. 2015. Heterogeneity and Bipotency of Astroglial-Like Cerebellar Progenitors along the Interneuron and Glial Lineages. J Neurosci, 35, 7388–402. 10.1523/jneurosci.5255-14.2015

Person, A. L. & Raman, I. M. 2011. Purkinje neuron synchrony elicits time-locked spiking in the cerebellar nuclei. Nature, 481, 502–5. 10.1038/nature10732

Renier, N., Wu, Z., Simon, D. J., Yang, J., Ariel, P. & Tessier-Lavigne, M. 2014. iDISCO: a simple, rapid method to immunolabel large tissue samples for volume imaging. Cell, 159, 896–910. 10.1016/j.cell.2014.10.010

Schwartz, P. M., Borghesani, P. R., Levy, R. L., Pomeroy, S. L. & Segal, R. A. 1997. Abnormal cerebellar development and foliation in BDNF−/− mice reveals a role for neurotrophins in CNS patterning. Neuron, 19, 269–81.

Sekerkova, G., Ilijic, E. & Mugnaini, E. 2004. Time of origin of unipolar brush cells in the rat cerebellum as observed by prenatal bromodeoxyuridine labeling. Neuroscience, 127, 845–58. 10.1016/j.neuroscience.2004.05.050

Sgaier, S. K., Lao, Z., Villanueva, M. P., Berenshteyn, F., Stephen, D., Turnbull, R. K. & Joyner, A. L. 2007. Genetic subdivision of the tectum and cerebellum into functionally related regions based on differential sensitivity to engrailed proteins. Development, 134, 2325–35. 10.1242/dev.000620

Sgaier, S. K., Millet, S., Villanueva, M. P., Berenshteyn, F., Song, C. & Joyner, A. L. 2005. Morphogenetic and cellular movements that shape the mouse cerebellum; insights from genetic fate mapping. Neuron, 45, 27–40. 10.1016/j.neuron.2004.12.021

Sillitoe, R. V., Gopal, N. & Joyner, A. L. 2009. Embryonic origins of ZebrinII parasagittal stripes and establishment of topographic Purkinje cell projections. Neuroscience, 162, 574–88. 10.1016/j.neuroscience.2008.12.025

Sillitoe, R. V. & Joyner, A. L. 2007. Morphology, molecular codes, and circuitry produce the three-dimensional complexity of the cerebellum. Annu Rev Cell Dev Biol, 23, 549–77. 10.1146/annurev.cellbio.23.090506.123237

Sonnier, L., Le Pen, G., Hartmann, A., Bizot, J. C., Trovero, F., Krebs, M. O. & Prochiantz, A. 2007. Progressive loss of dopaminergic neurons in the ventral midbrain of adult mice heterozygote for Engrailed1. J Neurosci, 27, 1063–71. 10.1523/jneurosci.4583-06.2007

Sudarov, A., Turnbull, R. K., Kim, E. J., Lebel-Potter, M., Guillemot, F. & Joyner, A. L. 2011. Ascl1 genetics reveals insights into cerebellum local circuit assembly. J Neurosci, 31, 11055–69. 10.1523/JNEUROSCI.0479-11.2011

Tan, I. L., Wojcinski, A., Rallapalli, H., Lao, Z., Sanghrajka, R. M., Stephen, D., Volkova, E., Korshunov, A., Remke, M., Taylor, M. D., Turnbull, D. H. & Joyner, A. L. 2018. Lateral cerebellum is preferentially sensitive to high sonic hedgehog signaling and medulloblastoma formation. Proc Natl Acad Sci USA. 10.1073/pnas.1717815115

Walberg, F. & Dietrichs, E. 1988. The interconnection between the vestibular nuclei and the nodulus: a study of reciprocity. Brain Res, 449, 47–53.

Wang, V. Y., Rose, M. F. & Zoghbi, H. Y. 2005. Math1 expression redefines the rhombic lip derivatives and reveals novel lineages within the brainstem and cerebellum. Neuron, 48, 31–43. 10.1016/j.neuron.2005.08.024

Wilson, S. L., Kalinovsky, A., Orvis, G. D. & Joyner, A. L. 2011. Spatially restricted and developmentally dynamic expression of engrailed genes in multiple cerebellar cell types. Cerebellum, 10, 356–72. 10.1007/s12311-011-0254-5

Wojcinski, A., Lawton, A. K., Bayin, N. S., Lao, Z., Stephen, D. N. & Joyner, A. L. 2017. Cerebellar granule cell replenishment postinjury by adaptive reprogramming of Nestin(+) progenitors. Nat Neurosci, 20, 1361–1370. 10.1038/nn.4621

Zhang, X. M., Ng, A. H., Tanner, J. A., Wu, W. T., Copeland, N. G., Jenkins, N. A. & Huang, J. D. 2004. Highly restricted expression of Cre recombinase in cerebellar Purkinje cells. Genesis, 40, 45–51.

Zhang, X. Y., Wang, J. J. & Zhu, J. N. 2016. Cerebellar fastigial nucleus: from anatomic construction to physiological functions. Cerebellum Ataxias, 3, 9. 10.1186/s40673-016-0047-1

