## Supplementary files for "Cerebellar nuclei neurons dictate cortical growth through developmental scaling of presynaptic Purkinje cells"

### Figure 1-figure supplement 1

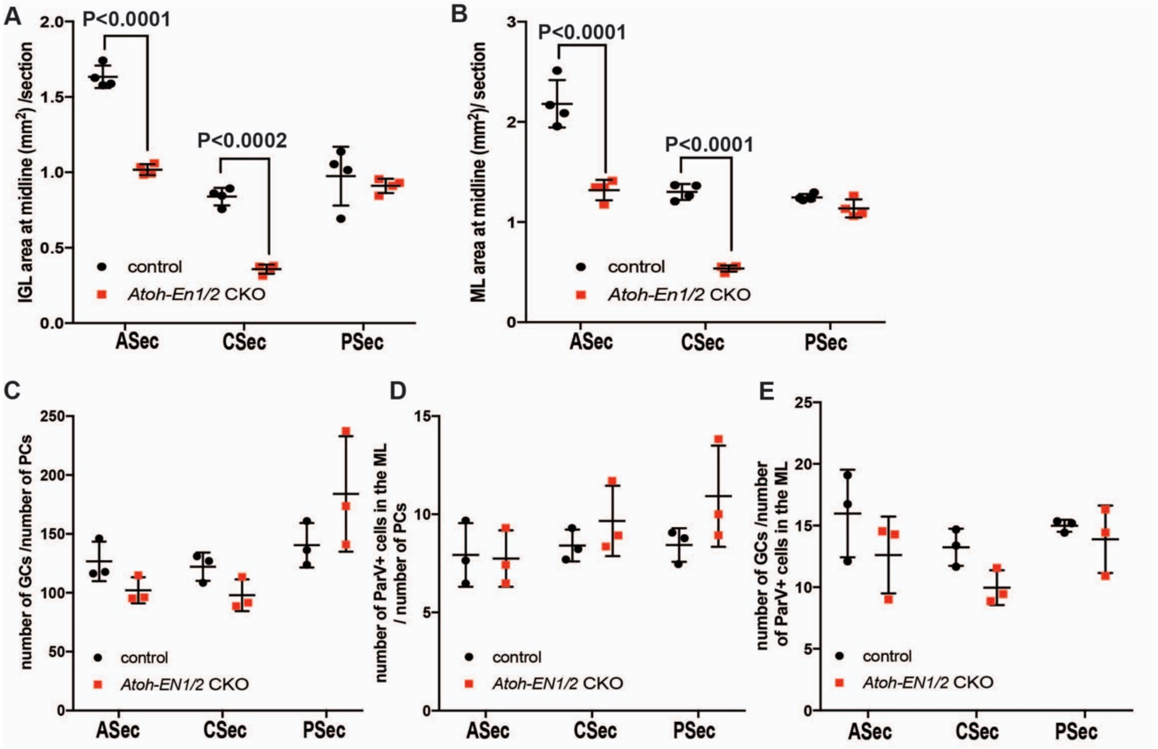


**Figure 1-figure supplement 1. IGL and ML area in the *Atoh-En1/2* CKO are reduced but the ratios of neurons in the cerebellar cortex remain similar to normal. A-B)** Quantifications of IGL (A) and ML (B) sector areas in *Atoh-En1/2* CKO animals compared to controls (n=4/condition, Two-way ANOVA, IGL: $F_{(1,9)}=98.8$, $P<0.0001$, ML: $F_{(1,9)}=278.3$, $P<0.0001$). **C-E)** Quantifications of the ratios of the numbers of granule cells (GCs) to PCs (C), ParV+ cells in the in the molecular layer (ML) to number of PCs (D) and the number of GCs to ParV+ cells in the ML in *Atoh-En1/2* CKO animals compared to controls (n=3/condition). Significant *post hoc* comparisons are shown in the figure.

**Figure 1-figure supplement 2**

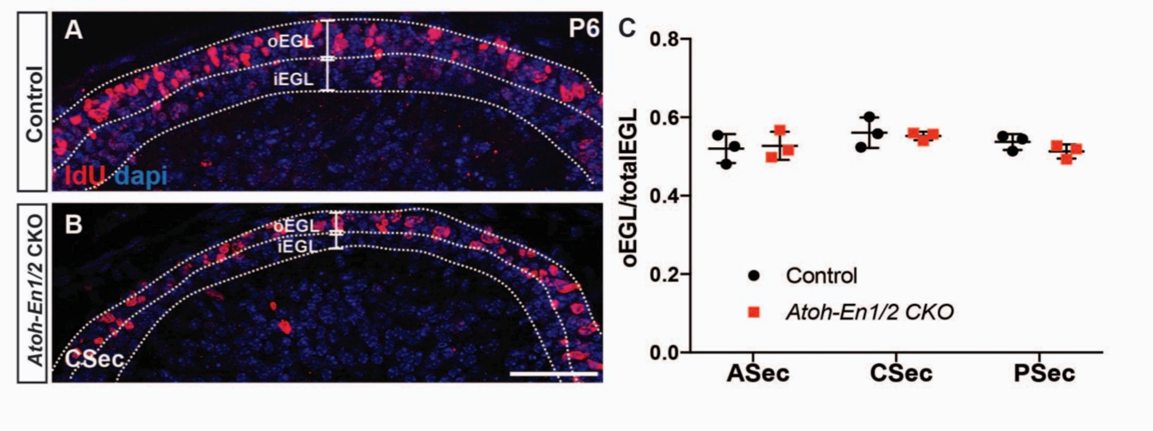


**Figure 1-figure supplement 2. Proportion of EGL that contains proliferating GCPs to total EGL area is preserved in *Atoh-En1/2* CKO animals. A-B)** Animals were injected with nucleotide analog IdU and sacrificed 1 hour later. Outer layer (oEGL) contains proliferating GCPs, whereas inner EGL (iEGL) contains differentiated granule cells. **C)** Quantification of oEGL to total EGL area shows that despite the smaller EGL at P6 in *Atoh-EN1/2 CKO* animals, the ratio of oEGL to total EGL is not changed (n=3/genotype, Two-way ANOVA, $F_{(1,6)}$=0.34, P=0.89).

**Figure 3-figure supplement 1**

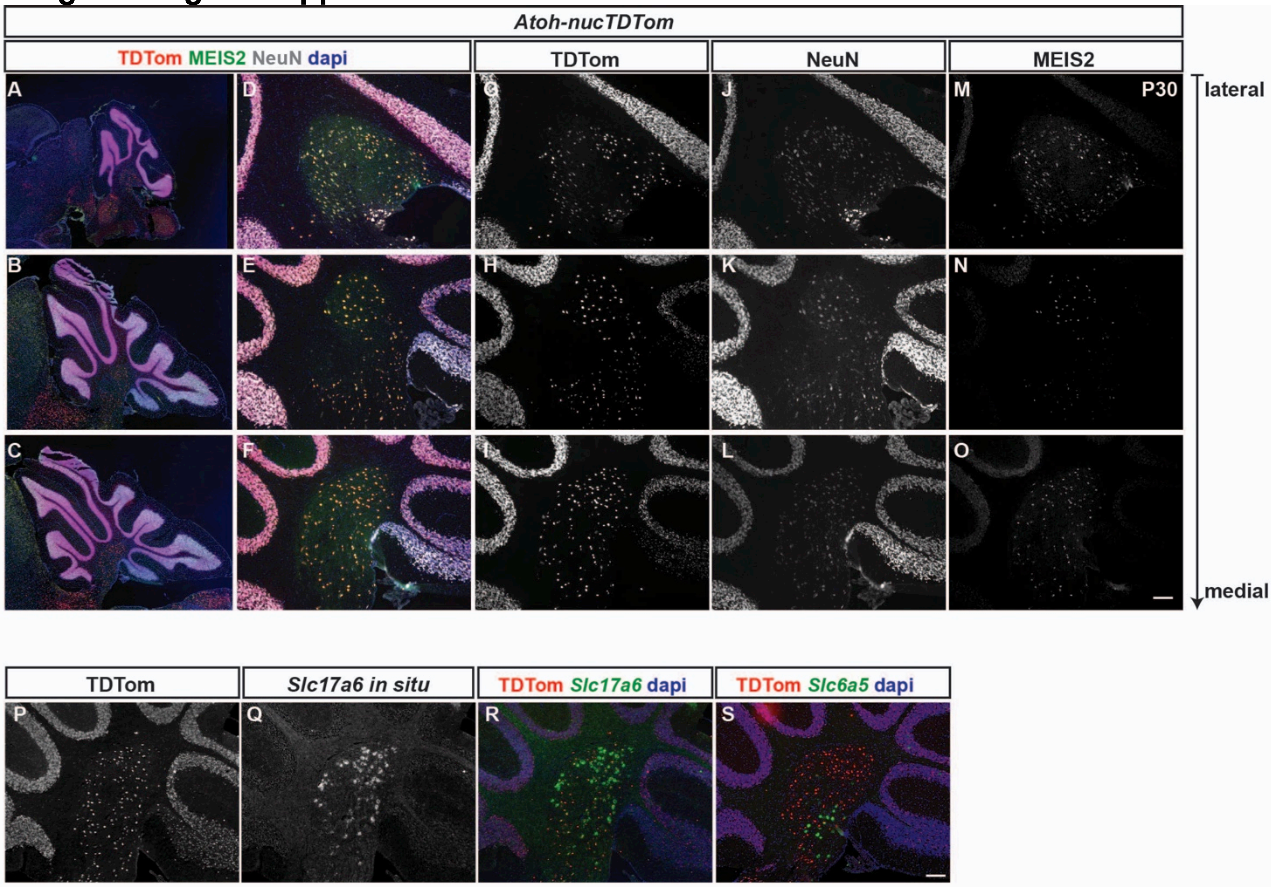


**Figure 3-figure supplement 1. *Atoh1-Cre* marked cells expressing TDTom express eCN markers (MEIS2 and *Slc17a6*) and a pan neuronal marker (NeuN) but not a glycinergic neuron marker *Slc6a5*. A-C)** Low power images of sagittal sections from the lateral (top) to medial (bottom) cerebellum of an *Atoh-TDTom* mouse are shown to indicate the level from which the images in (D-O) were taken. **D-O)** Co-labeling of TDTom, MEIS2, and NeuN showing that all TDTom+ cells (G-I) express NeuN (J-L) and the majority express MEIS2 (M-O) with the least at intermediate levels. **P-R)** *In situ* hybridization of *Slc17a6* (*Vglut2*) showing that the majority of TDTom+ cells express *Slc17a6*, although the levels of expression vary between cells. **S)** *In situ* hybridization analysis showing that only rare TDTom+ cells (<1 cell per section) express *Slc6a5* RNA (glycinergic neuron marker). *In situ* hybridization images were pseudocolored in Green.

Scale bars: 100 μm.

**Figure 3-figure supplement 2**

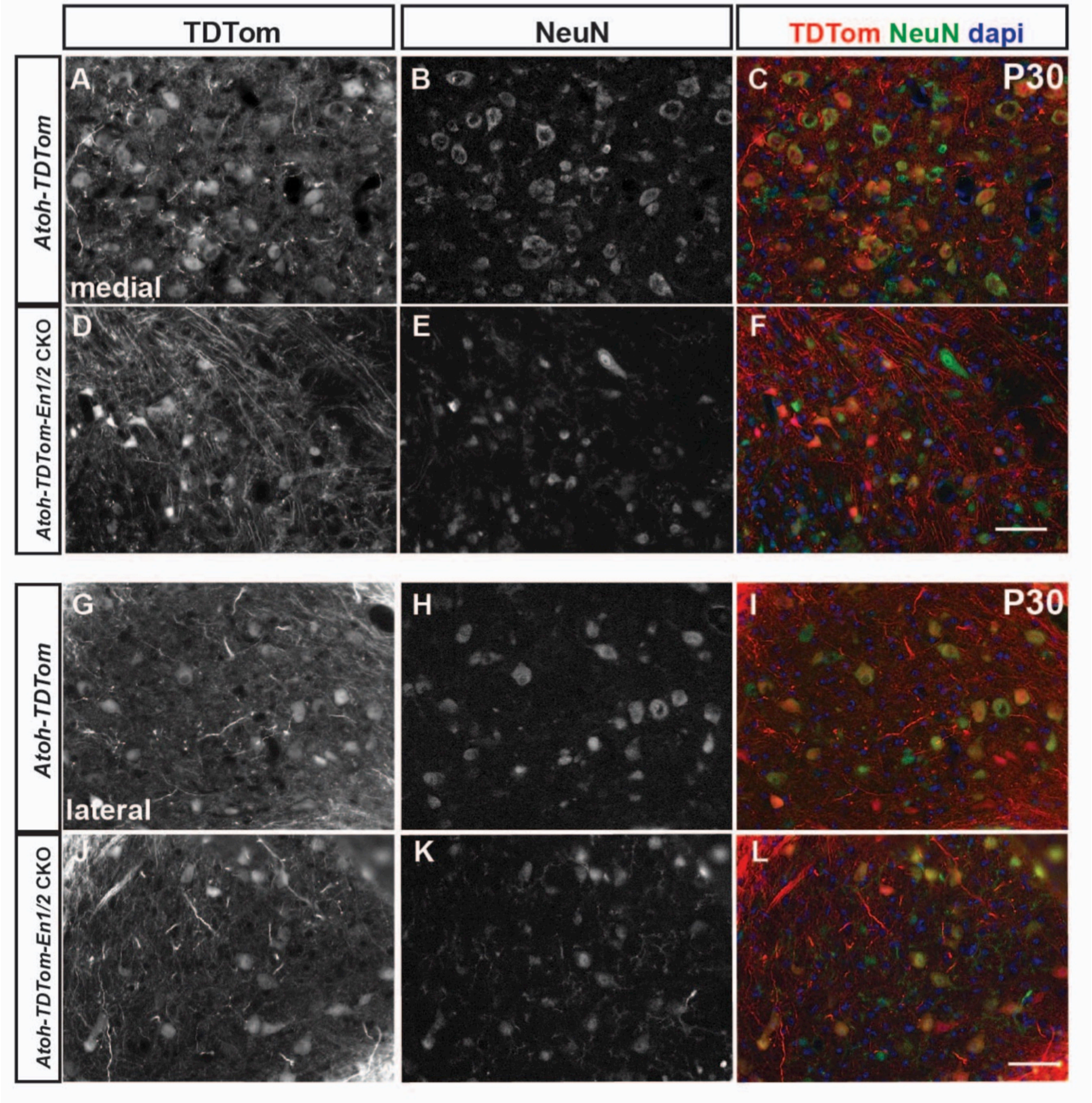


**Figure 3-figure supplement 2. Adult *Atoh-TDTom-En1/2* CKOs exhibit loss of medial eCN cells. A-L)** Immunofluorescent analysis of sagittal sections of eCN cells (NeuN+) marked by *Atoh1-Cre* (TDTom+) showing that the number of eCN cells is reduced in *Atoh-TDTom-En1/2* CKOs (D-F, J-L) compared *Atoh-TDTom* controls (A-C, G-I) in medial (A-F) but not lateral sections (G-L). Scale bars: 100 μm.

**Figure 4-figure supplement 1**

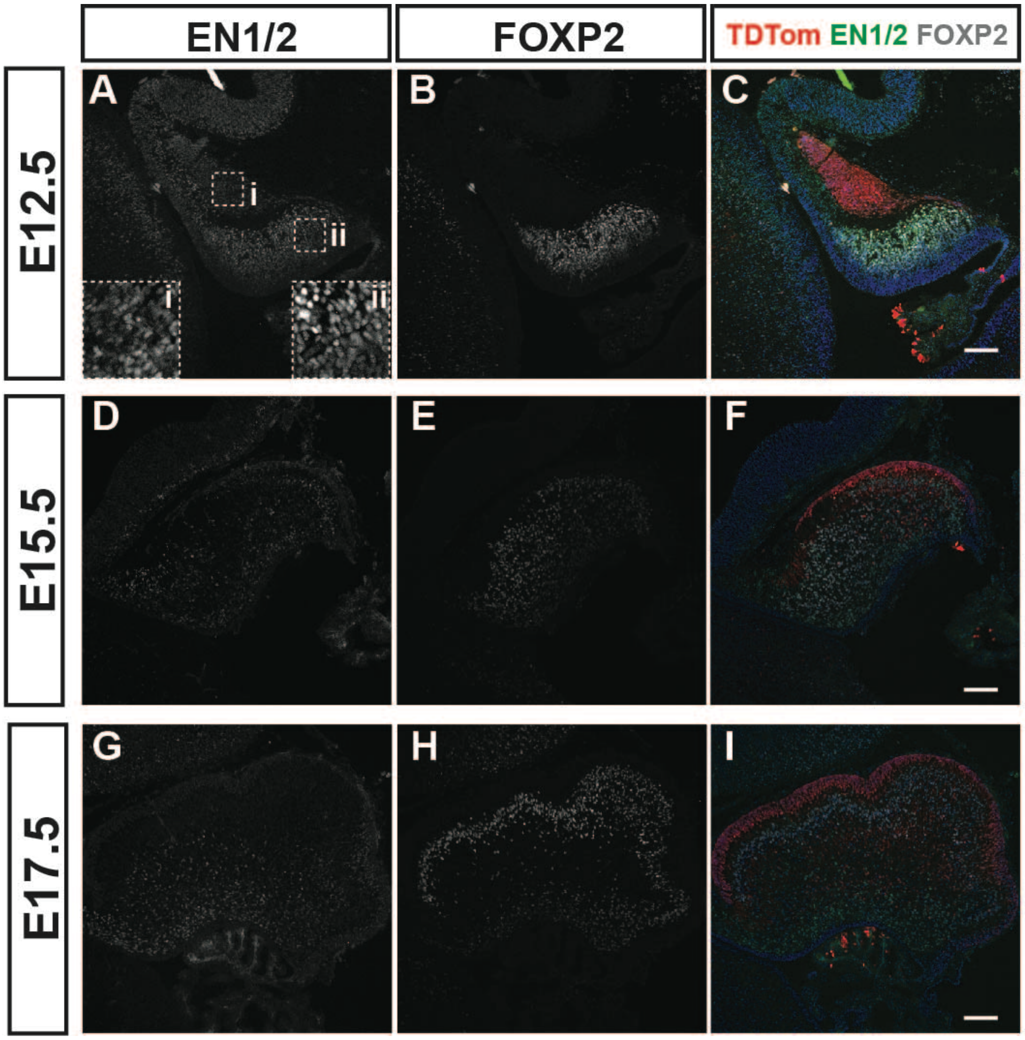


**Figure 4-figure supplement 1. Dynamic embryonic expression of EN1/2 in PCs during development. A-I)** Immunofluorescent analysis of midline sagittal sections from *Atoh-TDTom* animals showing co-expression of EN1/2 in TdTom+ cells (EGL and eCN) and in FOXP2+ PCs mainly at E12.5 (A-C), with decreasing expression at E15.5 (D-F) and E17.5 (G-I). Scale bars: 100 μm.

**Figure 5-figure supplement 1**

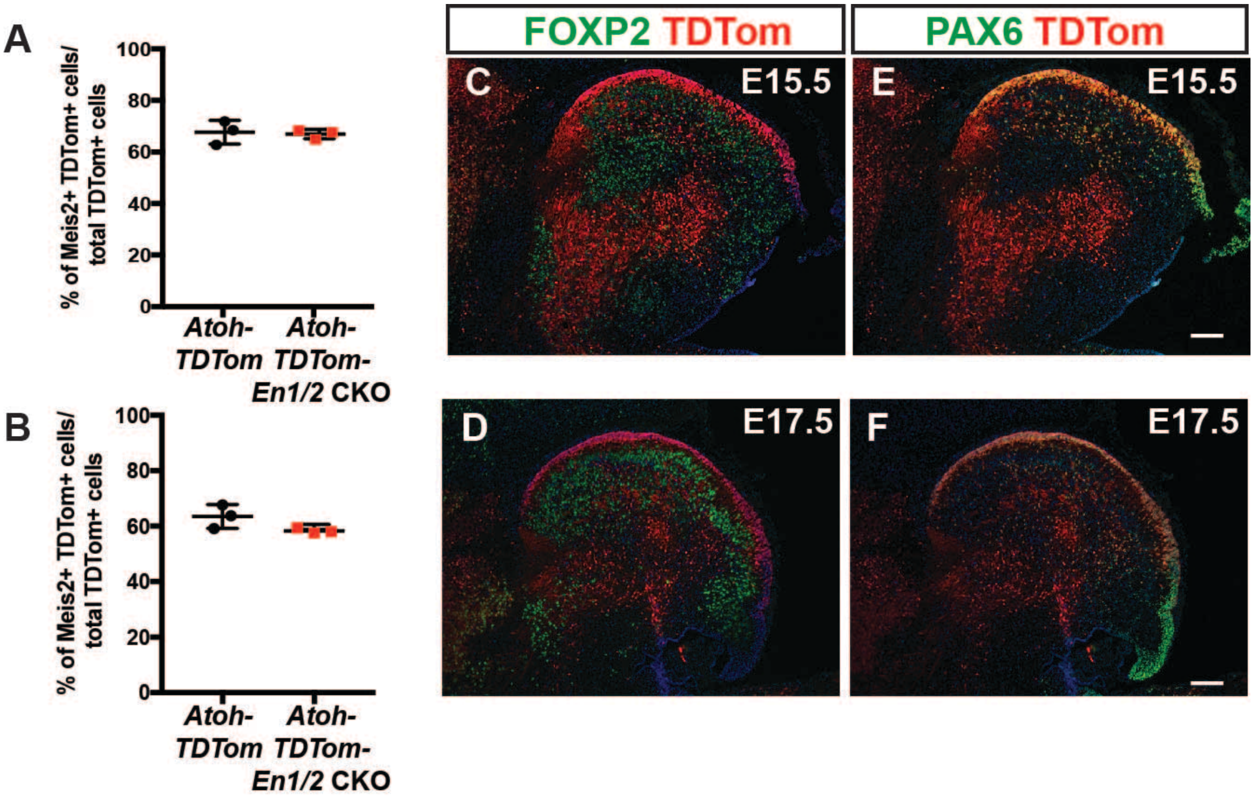


**Figure 5-figure supplement 1. *Atoh-Cre* marks the rhombic lip lineage but not Purkinje cells. A-B)** Quantification of the percentage of *Atoh1-Cre*-derived cells (TDTom+) in *Atoh-TDTom* mice that express MEIS2 showing no significant differences between *Atoh-En1/2* CKO mutants and controls at E15.5 (A) and E17.5 (B). **C-F)** Immunofluorescent analysis showing that TDTom+ cells in the EGL and a few in the cortex express PAX6 (marker for the granule cell lineage) at E15.5 (C) and E17.5 (D), and do not express the PC maker FOXP2+ (E-F) at either age. Scale bars: 100 µm.

**Figure 5-figure supplement 2**

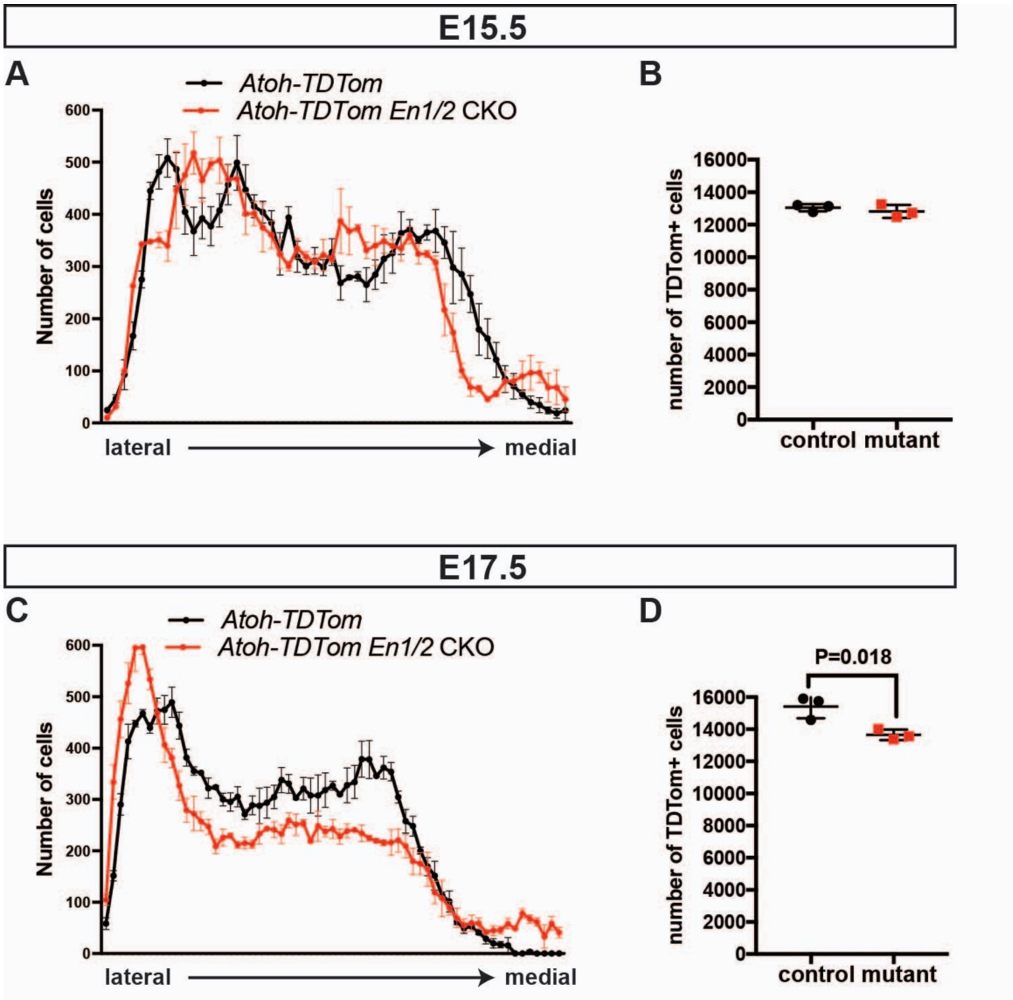


**Figure 5-figure supplement 2. EN1/2 loss leads to a reduction in eCN cells that are marked by *Atoh1-Cre*. A-B)** Quantification of the average number of TDTom+ eCN cells in each sagittal section along the medial-lateral axis (A) and total eCN numbers (B) in *Atoh-TDTom* and *Atoh-TDTom-En1/2* CKOs at E15.5 (n=3 for each genotype, Student's t-test, P=0.43). **C-D)** Quantification of the average number of TDTom+ eCN cells in *Atoh-TDTom* and *Atoh-TDTom-En1/2* CKOs at E17.5 (n=3 for each genotype, Student's t-test).

**Figure 5-figure supplement 3**

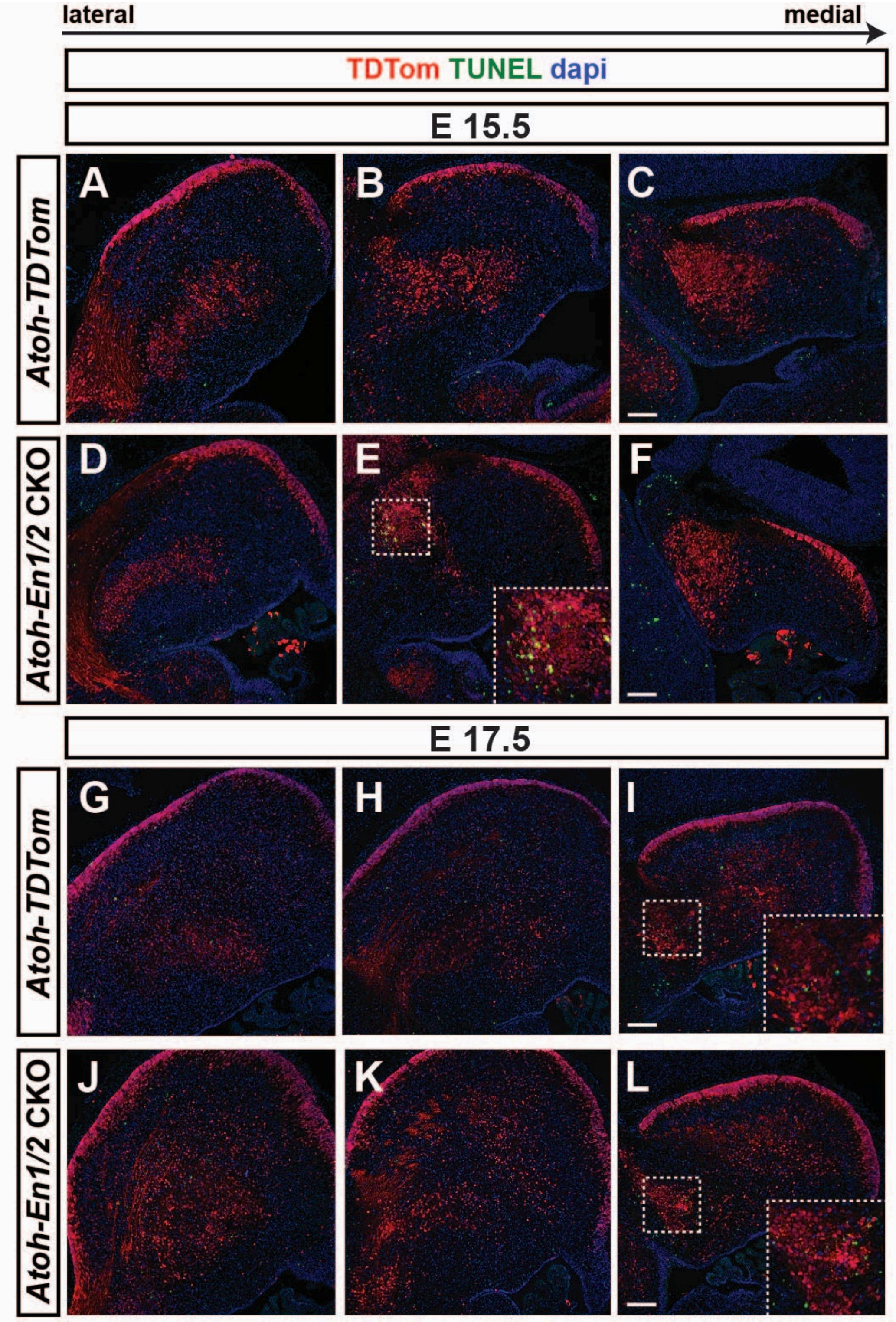

**Figure 5-figure supplement 3. Apoptosis is detected in the CN region following loss of EN1/2. A-L)** Immunofluorescent analysis of TDTom and TUNEL on sagittal sections from *Atoh-TDTom* (A-C, G-I) and *Atoh-TDTom-En1/2* CKO (D-F, J-L) animals showing that apoptosis is observed in the intermediate CN of CKO cerebella at E15.5 (A-F) and in more medial CN of CKO cerebella at E17.5 (G-L). Scale bars: 100 µm.

**Figure 7-figure supplement 1**

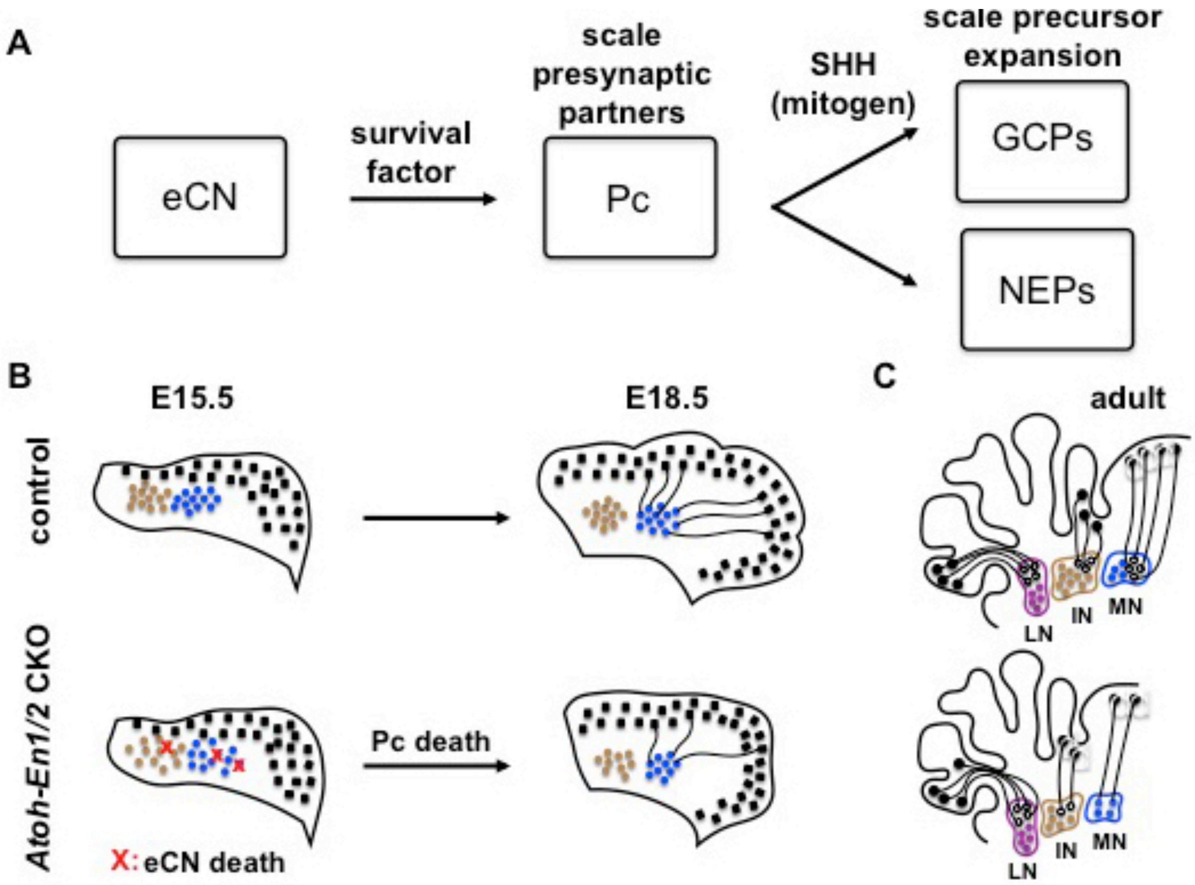


**Figure 7-figure supplement 2. Schematic representation of our findings and model.** The number of excitatory cerebellar nuclei neurons (eCN) scales the number of Purkinje cells (PCs) by secreting a survival factor for PCs, and granule cell, interneuron, and astrocyte numbers are expanded proportional to the number of PCs through expression of SHH.

**Figure 7-figure supplement 2**

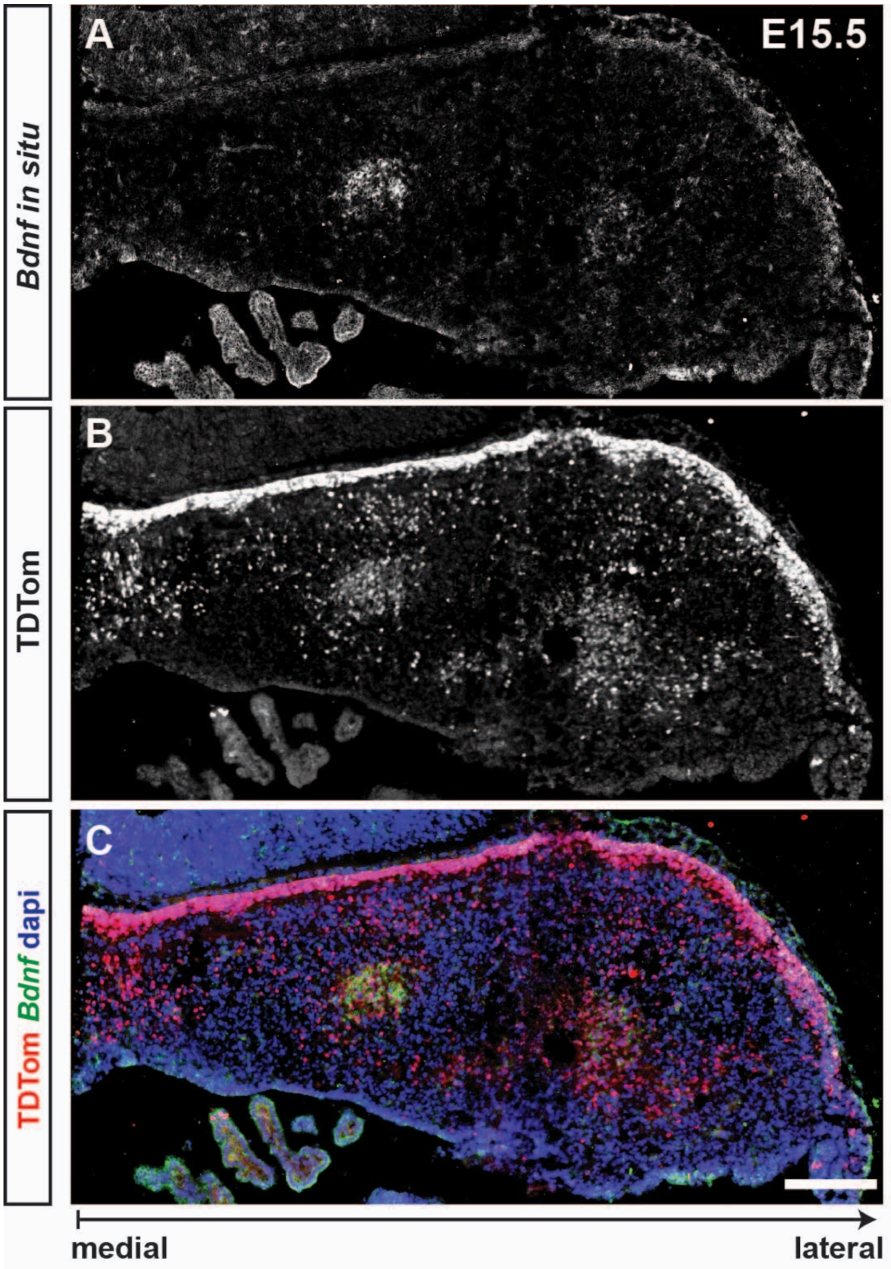


**Figure 7-figure supplement 1. RNA *in situ* hybridization of *Bdnf* shows expression in a subpopulation of eCN at E15.5. A-C)** *In situ* hybridization of analysis of *Bdnf* and antibody staining for TDTom protein on a coronal section of an E15.5 *Atoh-TDTom*

embryo showing *Bdnf* expression in a subpopulation of TDTom+ cells, likely the precursors of medial and some intermediate eCN*.* *In situ* hybridization images were pseudocolored in Green in (C). Scale bar: 200 µm

**Supplementary movie 1. 3D reconstruction of the stereology analysis done on a half cerebellum of a control animal at P30.** Green: medial CN; yellow: Intermediate CN; red: Lateral CN

**Supplementary movie 2. 3D reconstruction of the stereology analysis done on a half cerebellum of an *Atoh-EN1/2* CKO animal at P30.** Green: medial CN; yellow: Intermediate CM; red: Lateral CN

### Source Data_1

| Source Data_1 | | | | |
|---|---|---|---|---|
| **Figure** | **Test performed** | **P-Value** | **Multiple comparisons** | |
| Fig. 1A | Unpaired t-test | t(6)=21.99 | N/A | |
| Fig. 1C | Two-way ANOVA | $F_{(1, 9)}$=398.277, P<0.0001 | ASec | P<0.0001 |
| | | | CSec | P<0.0001 |
| | | | PSec | 0.1498 |
| Fig.1D | Two-way ANOVA | $F_{(1, 9)}$=0.2269, P=0.64 | ASec | 0.69 |
| | | | CSec | 0.9661 |
| | | | PSec | 0.9955 |
| Fig. 1J | Two-way ANOVA | $F_{(1, 15)}$=72.52, P<0.0001 | ASec | P<0.0001 |
| | | | CSec | P<0.0001 |
| | | | PSec | 0.3064 |
| Fig. 1K | Two-way ANOVA | $F_{(1, 15)}$=0.2583, P=0.6187 | ASec | 0.9484 |
| | | | CSec | 0.8998 |
| | | | PSec | 0.8512 |
| Fig. 1L | Two-way ANOVA | $F_{(1,9)}$=0.8772, P=0.3734 | ASec | 0.3167 |
| | | | CSec | 0.7412 |
| | | | PSec | 0.6820 |
| Fig.1M | Two-way ANOVA | $F_{(1,9)}$=28.4, P<0.0005 | ASec | 0.03 |
| | | | CSec | 0.003 |
| | | | PSec | 0.5937 |
| Fig. 2A | Student's t-test | d1 run1: t(8)=0.2373 P=0.8184<br>d1 run2: t(8)=0.7541 P=0.4724<br>d1 run3: t(8)=0.5988 P=0.5658<br>d2 run1: t(8)=0.8093 P=0.4417<br>d2 run2: t(8)=2.033 P=0.1109<br>d2 run3: t(8)=1.422 P=0.2439<br>d3 run1: t(8)=2.135 P=0.07<br>d3 run2: t(8)=2.86 P=0.03<br>d3 run3: t(8)=2.651 P=0.034 | N/A | |
| Fig. 2B | Student's t-test | t(15)=2.683 P=0.02 | N/A | |
| Fig. 2C | Student's t- | Stride: t(25)=4.198 | N/A | |

| | | | | |
|---|---|---|---|---|
| | test | P=0.0003<br>Sway: t(25)=2.398<br>P=0.03<br>Stance: t(25)=2.219<br>P=0.01 | | |
| Fig. 2D | Student’s t-test | t(15)=3.656 P=0.002 | N/A | |
| Fig. 3J | Two-way ANOVA | F (1, 12) = 32.29<br>P=0.0001 | ASec | 0.0058 |
| | | | CSec | 0.0001 |
| | | | PSec | 0.8003 |
| Fig. 3M | Student’s t-test | t(5)=6.397 P=0.014 | N/A | |
| Fig. 4E | Student’s t-test | t(4)=3.085 P=0.04 | N/A | |
| Fig. 5B | Student’s t-test | t(4)=0.1561 P=0.88 | N/A | |
| Fig. 5J | Student’s t-test | t(4)=8.514 P=0.001 | N/A | |
| Fig. 6C | Student’s t-test | t(4)=0.4378 P=0.6841 | N/A | |
| Fig. 6D | Student’s t-test | t(4)=1.418 P=0.2293 | N/A | |
| Fig. 6G | Student’s t-test | t(4)=8.93 P=0.0009 | N/A | |
| Fig. 6H | Student’s t-test | t(4)=6.559 P=0.0028 | N/A | |
| Fig. 7C | Student’s t-test | aMN: t(4)=4.492<br>P=0.0109<br>pMN: t(4)=4.412<br>P=0.0116 | N/A | |
| Fig1_supplement 1A | Two-way ANOVA | $F_{(1,9)}$=98.8, P<0.0001 | ASec | 0.0001 |
| | | | CSec | 0.0002 |
| | | | PSec | 0.7396 |
| Fig1_supplement 1B | Two-way ANOVA | $F_{(1,9)}$=278.3, P<0.0001 | ASec | 0.0001 |
| | | | CSec | 0.0001 |
| | | | PSec | 0.2706 |
| Fig1_supplement 2C | Two-way ANOVA | $F_{(1,6)}$=0.34, P=0.89 | ASec | 0.9935 |
| | | | CSec | 0.9887 |
| | | | PSec | 0.8213 |
| Fig 5_supplement 1A | Student’s t-test | t(4)=0.2472 P=0.8169 | N/A | |
| Fig 5_supplement 1B | Student’s t-test | t(4)=2.048 P=0.11 | N/A | |

| | | | |
|---|---|---|---|
| Fig 5_supplement 2B | Student’s t-test | t(4)=0.869 P=0.4339 | N/A |
| Fig 5_supplement 2E | Student’s t-test | t(4)=3.83 P=0.018 | N/A |
